# The distribution of distances to the edge of species coexistence

**DOI:** 10.1101/2024.01.21.575550

**Authors:** Mario Desallais, Michel Loreau, Jean-François Arnoldi

## Abstract

In Lotka-Volterra community models, given a set of biotic interactions, recent approaches have analysed the probability of finding a set of species intrinsic growth rates (representing intraspecific demographic features) that will allow coexistence. Several metrics have been used to quantify the fragility of coexistence in the face of variations in those intrinsic growth rates (representing environmental perturbations), thus probing a notion of ‘distance’ to the edge of coexistence of the community. Here, for any set of interacting species, we derive an analytical expression for the whole distribution of distances to the edge of their coexistence. Remarkably, this distribution is entirely driven by (at most) two characteristic distances that can be directly computed from the matrix of species interactions. We illustrate on data from experimental plant communities that our results offer new ways to study the contextual role of species in maintaining coexistence, and allow us to quantify the extent to which intraspecific features and biotic interactions combine favorably (making coexistence more robust than expected), or unfavourably (making coexistence less robust than expected). Our work synthesizes different study of coexistence and proposes new, easily calculable metrics to enrich research on community persistence in the face of environmental disturbances.

## Introduction

Understanding why and how species coexist is a central question in community ecology (Armstrong and McGehee, 1976; Chesson, 2000; Hastings, 1980; Hutchinson, 1961). Many studies have focused on what makes coexistence possible, and in particular on the role of the network of interactions between species (Abrams, 1984; Abrams et al., 2003; Brose et al., 2006; Otto et al., 2007; Williams, 2008). In the context of Lotka-Volterra models (the simplest mathematical representations of the population dynamics of interacting species), to quantify the role played by biotic interactions in species coexistence, a recent and growing body of theoretical work proposes to study the volume of a community’s so called ‘feasibility domain’ (Rohr et al., 2014, 2016; Saavedra et al., 2017; Song et al., 2018b). Given the set of biotic interactions between species, this feasibility domain is defined as the range of species intrinsic features (thought to reflect abiotic conditions that do not depend on the presence of the other species considered, such as intrinsic growth rates or carrying capacities) that allow species to coexist (Fig. 1). The idea here is that the larger this domain, the more likely a community is to withstand environmental disturbances while maintaining coexistence (Bartomeus et al., 2021; Song et al., 2018a).

**Figure 1.**
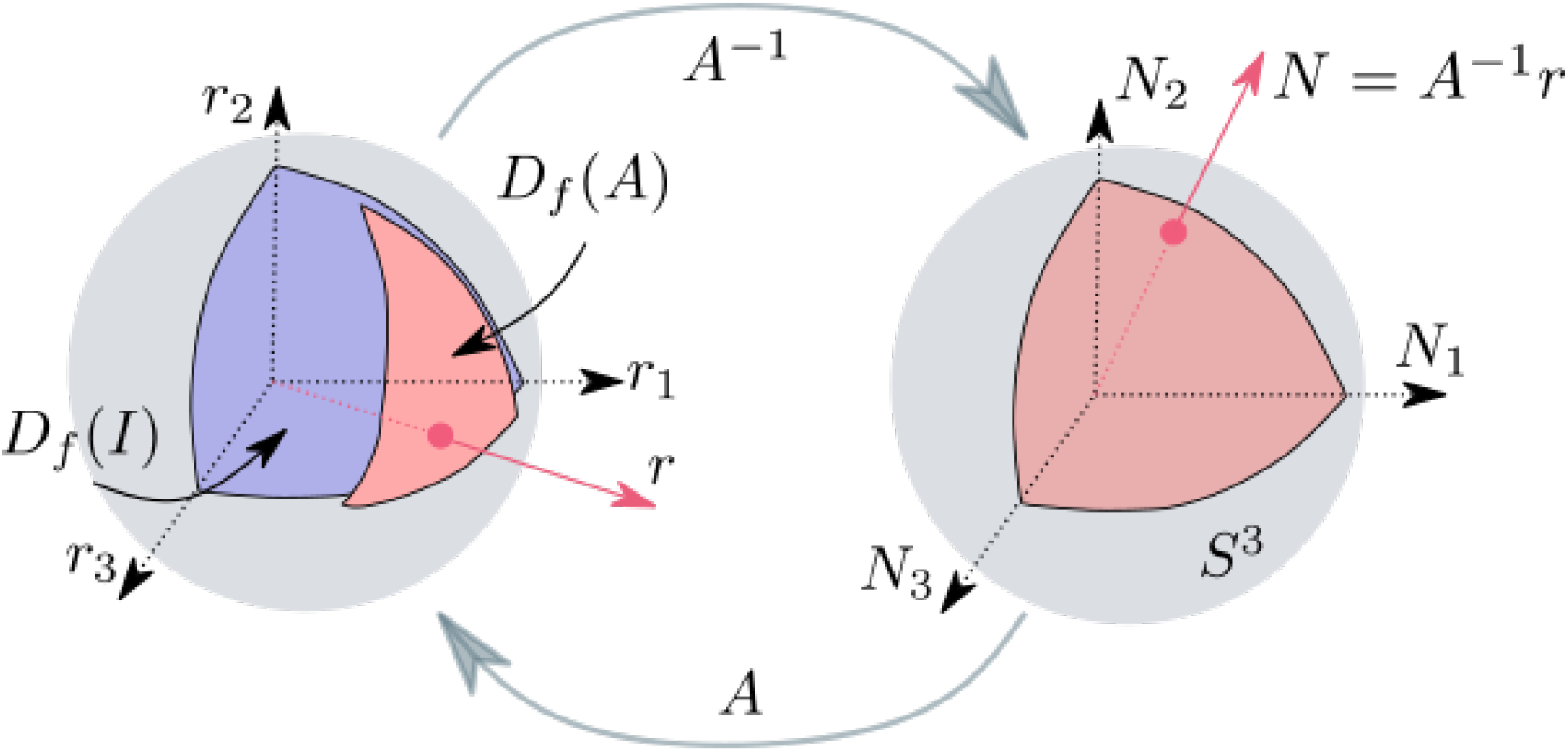
The feasibility domain *D*_*f*_ (*A*) (in light red on the left) is defined as the subset of growth rate directions that, given a pair-wise interaction matrix *A*, allows coexistence between all species. It is the intersection of the sphere with the image in r-space (via the matrix *A*) of the positive quadrant in N-space (shown on the right). The shape and volume of the feasibility domain corresponds to the shape and volume of the light red surface on the left. The probability of feasibility ℙ (*r* ∈ *D*_*f*_) is the ratio between the volume of *D*_*f*_ and the volume of the unit sphere.

However, the fact that a large set of conditions allows coexistence does not necessarily mean that coexistence is robust to environmental change. A thin elongated feasibility domain could have a large volume, yet only contain fragile coexistence states, vulnerable to small changes in abiotic conditions. This observation reflects the tenuous distinction between two seemingly equivalent questions: “how likely will species coexist?”, whose answer, in L-V models, corresponds to the size of the feasibility domain, and “If species do coexist, how fragile will this coexistence be?”. This difference between raw and conditional probabilities of coexistence has led to the emergence of shape metrics of feasibility domains (Allen-Perkins et al., 2023; Grilli et al., 2017; Saavedra et al., 2017).

In line with these recent approaches, the aim of our study is to expand on the study of feasibility by proposing an explicit mathematical relationship between the robustness of coexistence in the face of environmental disturbances, and the shape and size of a feasibility domain. To do so, we model ecological perturbations as long term changes of species intrinsic features (such as their growth rates or carrying capacities) and define, for any realized coexistence state, a notion of distance to the edge of coexistence. This distance is the minimal environmental perturbation intensity *z* able to lead at least one species to extinction. Our goal is to determine, amongst all coexistence states, the proportion *p*(*z*) that lie within distance *z* from the edge of feasibility. For a given feasibility domain, this function *z* ↦ *p*(*z*) describes the distribution of distances to its edges, thus characterizing both the size and shape of the domain. If the function *p*(*z*) rapidly reaches 1 as *z* grows this means that coexistence is typically fragile. The (cumulative) function *p*(*z*) quantifies the interrelation between species growth rates and their interactions. For instance, if in a given state, *p*(*z*) is close to 1, this means that in this environment, the set of species intrinsic growth rates and the set of their biotic interactions combine favourably. Our mathematical analysis will reveal the essential features of the function *p*(*z*) that can be directly computed from the matrix of biotic interactions.

As we hinted above, our description of the distribution of distances to the edge of coexistence is in line with recent work by Allen-Perkins et al. (2023). Using a similar logic to study the asymmetry of the feasibility domain (but different analytical calculations) these authors introduced different metrics related to the robustness of coexistence of the community. Remarkably, they used one of these metrics, the so-called “probability of exclusion”, to characterize species vulnerability in grasslands, showing that theoretical predictions based on the shape of the feasibility domain are consistent with observed population dynamics. In a similar vein we show here how to use features of the function *p*(*z*) to study the relative vulnerability of species. The idea is to address the biotic role played by each species in the robustness of coexistence, in the context of the community to which it belongs.

We apply our methods to simulated ecological communities, either drawing parameters at random (See appendix B) or inferring them from experimental plant community experiments (Van Ruijven and Berendse, 2009). The results (in line with Allen-Perkins et al. (2023)) confirm the link between the coexistence measures we derive from our work and the actual persistence of species through time in a changing environment. Applied to experimental plant community data, our analysis reveals the role played by the various plant species in maintaining coexistence, which we relate to the degree of facilitation or competition experienced by each. We also quantify the adequacy, in terms of coexistence, between biotic and abiotic conditions in those plant communities. Our work constitutes a proof of concept, demonstrating a theoretical method for future experiments aimed at characterizing a particular type of environment and how well it matches a particular assemblage of species in terms of maintaining coexistence.

### The feasibility domain

Consider a community of *S* species. Let *N*_*i*_ define the abundance of species *i* and *r*_*i*_ its intrinsic growth rate (which could be negative if the species cannot establish on its own), which encodes the effect of the environment on the ability of the species to grow if it were alone (Coulson et al., 2017; Levins, 1968; Meszéna et al., 2006; Roughgarden, 1975). The central object of study of feasibility is the matrix *A* = (*A*_*ij*_) of pairwise biotic interactions between all *S* species in the community. *A*_*ij*_ encodes how a change in the abundance of species *j*, impacts the growth of species *i*. This can represent competition or facilitation depending on the sign of *A*_*ij*_. The diagonal terms *A*_*ii*_ represent intraspecific competition, and will be assumed non-zero in our analysis. The generalized Lotka-Volterra (L-V) model (Volterra, 1926) prescribes the population dynamics of all species as:

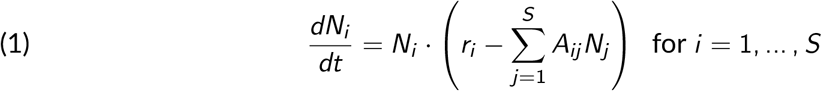

A growth rate vector *r* = (*r*_*i*_) is ‘feasible’ if the fixed point *N*^*^(*r*) = *A*^−1^*r* of the above model is strictly positive, meaning that *N*^*^(*r*)_*i*_ *>* 0 for all *i*. To guarantee the coexistence when the feasible equilibrium point is reached, we impose global stability of the system (Deng et al., 2022) by considering only D-stable interaction matrices (Grilli et al., 2017). To define the feasibility domain one has to assume that variations in growth rates are the result of a variation in abiotic conditions impacting the ability of species to grow on their own, but not their interactions (but see discussion). This abstraction leads to a definition of the feasibility domain associated with the interaction matrix *A* (Rohr et al., 2014): the set of growth rate vectors *D*_*f*_ (*A*) such that the equilibrium abundances are non-zero. However, in the L-V model, multiplying all growth rates by a constant does not change coexistence. Thus the feasibility domain has to be defined as a set of *directions*, isomorphic to a solid angle in the r-vector space (Ribando, 2006; Saavedra et al., 2017; Song et al., 2018b), so a convex subset of the sphere (Fig. 1):

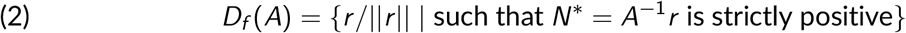

We can also think of the relative volume of the domain as the probability ℙ (*r* ∈ *D*_*f*_) of randomly drawing growth rates *r* which lead to positive abundances (Grilli et al., 2017). The random sampling must be thought of as uniform in the space of growth rate *directions*. Importantly, drawing each species’ growth rate *r*_*i*_ independently from a standard Gaussian distribution yields such a uniform sampling of growth rate directions. This remark, followed by the linear change of variables *A*^−1^ : *r* ↦ *N* then leads to the following formula:

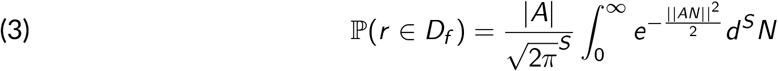

The probability ℙ (*r* ∈ *D*_*f*_) can therefore be computed as the cumulative distribution, evaluated at 0, of a normal distribution whose covariance matrix is determined by the interaction matrix *A* (this covariance matrix is (*A*^⊤^*A*)^−1^). In the absence of interactions ℙ (*r* ∈ *D*_*f*_) = 2^−*S*^. To focus on the effect of interactions it is thus convenient to define a ratio of probabilities (Saavedra et al., 2017):

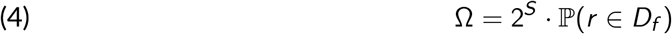

Ω corresponds to the effect of species interactions on the probability of coexistence and is equal to 1 in the non-interaction case.

These are well known results, and since their first introduction to ecology by (Rohr et al., 2014), have been applied to study the coexistence of many ecological systems. Yet the volume of the feasibility domain (also called structural stability) does not, a priori, tell us anything about the shape of the domain, nor how to relate its value to the probability that a given perturbation will push some species to extinction. Our goal in the next section is to provide such a connection.

### Distribution of distances from the edge of a triangle

If the community is made of three species (*S* = 3), the feasibility domain corresponds to a solid angle, a triangle drawn on a sphere (see Fig. 1). We thus start with a simplified analysis of regular triangles. This analogy allows us to gradually introduce the logic behind our geometrical approach (see Fig. 2). In this detour into simple trigonometry, which may seem removed from the initial ecological question, we will create a shape metric capable of encapsulating all the subtleties of shape differences between triangles (see Fig. 2).

**Figure 2.**
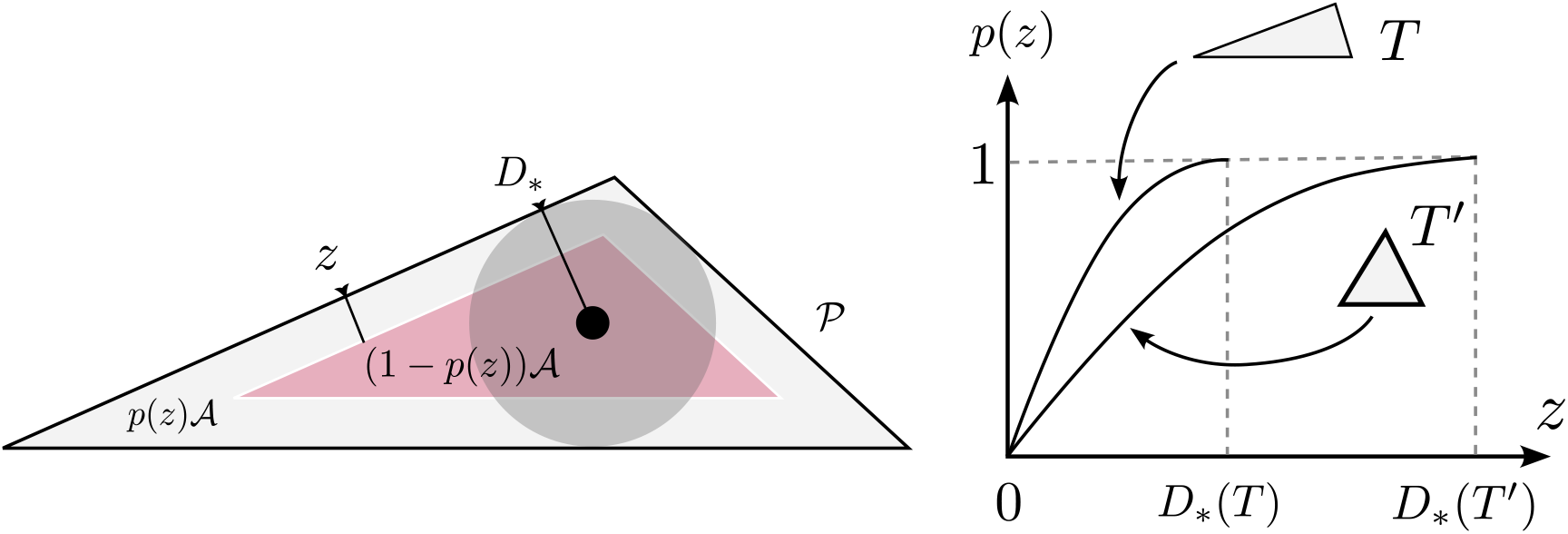
Left: Triangles are parameterized by their area 𝒜 and perimeter 𝒫. We are interested in the fraction *p*(*z*) of points that lie within a distance *z* from an edge. We can show that *p*(*z*) is fully parameterized by *D*_*_ = 2 𝒜*/𝒫*, the radius of the inscribed disc, whose center is equidistant to all edges of the triangle. Right: at a fixed area 𝒜, *D*_*_ grows as triangles become equilateral.

The probability of a point to be at a distance greater than *z* from one of the triangle’s edges corresponds to the relative area of the inscribed triangle whose own edges are exactly at a distance *z* from the boundaries of the original one (see the left panel of Fig. 2). Knowing 𝒜, the area of the original triangle, and 𝒜_*′*_, the area of the inscribed triangle, the proportion *p*(*z*) of points that lie within a distance *z* from an edge is thus 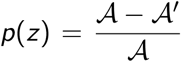. It is an entertaining exercise to show that

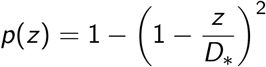

showing that *p*(*z*) is fully parameterized by a single number *D*_*_, which is the radius of the largest disc contained in the triangle (indeed *p*(*D*_*_) = 1). One can show that *D*_*_ = 2 𝒜*/ 𝒫* where 𝒫 is the perimeter of the original triangle. For a fixed area 𝒜, *D*_*_ is maximal for equilateral triangles (right panel of Fig. 2). This single distance measure *D*_*_ therefore allows us to quantitatively express differences in shape and size between triangles, and quantify, via the function *p*(*z*) which encodes the whole distribution of distances to the triangle’s edges.

In the next section we will generalize this geometrical ideas to feasibility domains, that are not simple triangles, and can be of any dimension (i.e. any number of species). The aim is now to derive a similar function *p*(*z*) applicable to ecological systems (L-V models).

### Distribution of distances from the edge of a feasibility domain

Following Allen-Perkins et al. (2023), Cenci et al. (2018a), and De Laender et al. (2023), we consider perturbations as changes in environmental conditions that occur on a long-time scale (so that a new equilibrium can be reached). Mathematically, we model a perturbation as a vector of variation *δr* of species intrinsic growth rates (i.e. whose components are the species-level variations *δr*_*i*_). Using the euclidean norm of vectors || *·* || we then measure the relative intensity of this perturbation as (we will see why below)

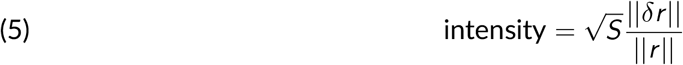

For any point *r* in a feasibility domain (so a feasible growth rate vector), we can measure its distance from the edge of the domain as the minimal perturbation intensity capable of leading at least one species to extinction. In the appendix we show that this distance can be directly computed as

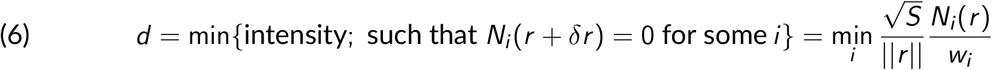

in the last term, for any species *i, w*_*i*_ is the euclidean norm of the corresponding row of the inverse interaction matrix, which encodes that species sensitivity to environmental perturbations, with *w*_*i*_ measuring its maximal sensitivity (thus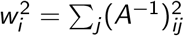). Our main result, illustrated in Fig. 3, is a simple formula for the distribution of such distances, in the form of a cumulative function *p*(*z*) = ℙ (*d* ≤ *z*), which mimics the one given in the previous section for standard triangles, and is entirely parameterized by two characteristic distances and species richness 𝒮:

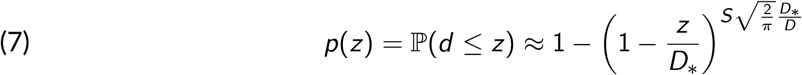

**Figure 3.**
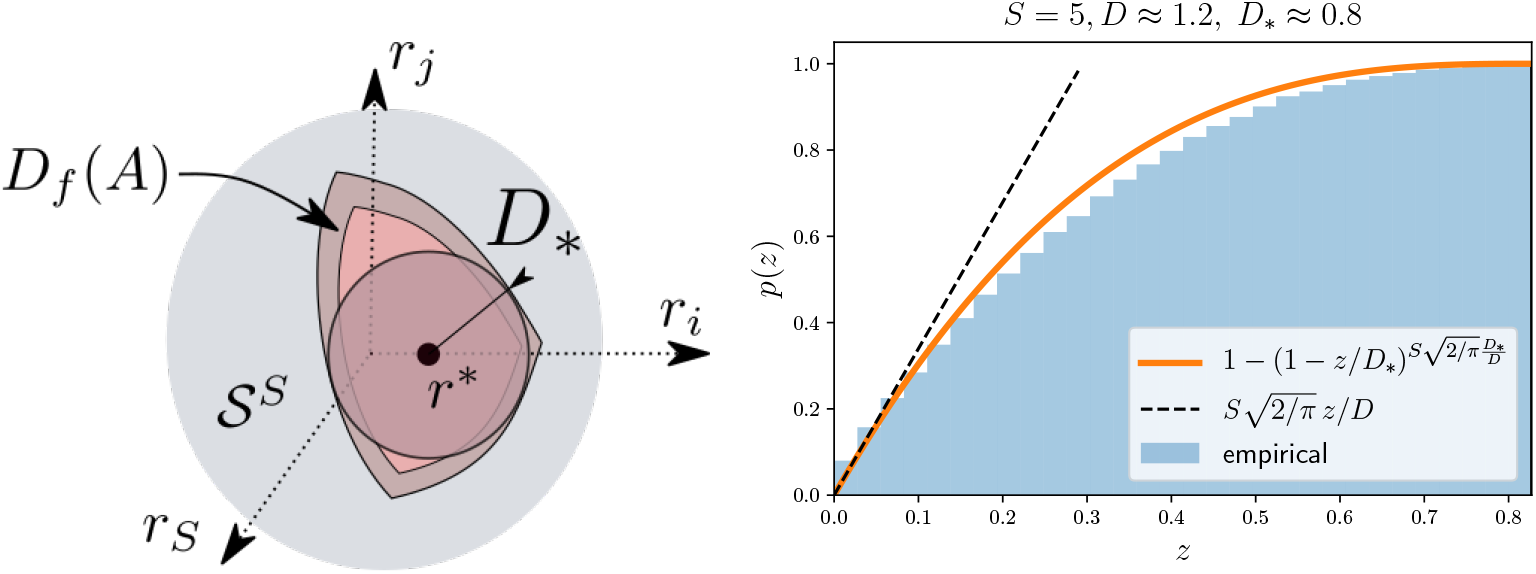
Empirical distribution of distances *p*(*z*) for a random interaction matrix of 𝒮 = 5 species, compared with its analytical approximation (equation 7 in orange line). To obtain the empirical distribution, we randomly sampled distances to the edge *z* and calculated, for each distance *z*, the fraction *p*(*z*) of coexistence states closest to the edge.

As for standard triangles, *D*_*_ represents the largest distance within the domain, associated with its incenter *r*^*^, also the most robust state of coexistence given the set of biotic interactions. Remarkably, we can deduce a simple formula for both *D*_*_ and *r*^*^. Indeed, in the appendix we show that, if *w* is the vector of maximal species sensitivities (i.e. whose components are the species-level values *w*_*i*_), then (8)

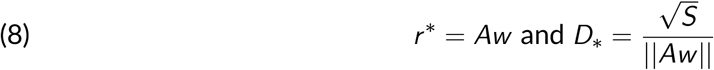

One can check that in the absence of interactions, and thus when *A* is diagonal, we have *D*_*_ = 1 (this is a consequence of our choice of normalisation of perturbation intensity). The formula for the distribution of distances differs from the one for triangles in that the maximal distance is not the only relevant distance, the one driving the behaviour at small *z* values, so near the edge of the domain, is in fact

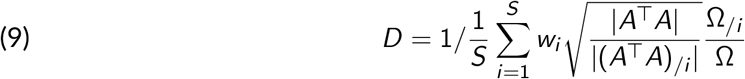

with *X*_*/i*_ notation meaning for any matrix *X*, the corresponding matrix without the *i* − *th* row and column, and Ω_*/i*_ is essentially the relative volume of the feasibility domain for the community without species *i* (but see the appendix for a more precise expression and derivation). The initial slope of *p*(*z*) is given by 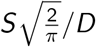 and determines the behavior of *p*(*z*) at small *z* values, so near the edge of the domain. We see that this slope explicitly grows with species richness. The latter behavior occurs because when there are many species present, it is ever more likely that one of them is close to local extinction. This diversity effect will tend to take a dominant part in shaping the function *p*(*z*). Geometrically speaking, this effect comes from the fact that in high dimensions, even very thin neighbourhoods of the edge of a closed object will cover a dominant fraction of the overall volume of that object. The expression for *D* and *D*_*_ clearly differ. Nonetheless those two distances are closely related and take very similar values, with *D* ≈ *D*_*_ for the vast majority of random interactions matrices that we generated, and even more so when considering empirically inferred matrices (See supplementary figure A2). Finally, we can connect the characteristic distances *D*_*_ and *D* with the relative volume of the domain Ω (from the first section). In supplementary figure A1 and in the mathematical appendix we explain why we may expect that, roughly speaking

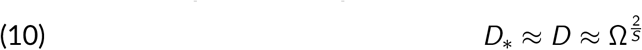

This approximate relationship, which must be understood as an equivalence of orders of magnitudes and not of precise values, taken together with equation 7 connects the size of the domain to its shape, and to the probability that a given perturbation can push species to extinction. In the appendices we expand on the latter point by considering randomly changing environments trough time. We show in simulations that the equivalent metrics of equation 10 do predict the duration of stable coexistence periods (See supplementary figure B1 in the “Persistence of species in simulated ecological system” appendix).

### Contextual species vulnerability

The above analyses of the distribution of distances to the edge of feasibility enable us to characterize the robustness of coexistence of an ecological community. We now take the analysis further to show that the characteristic distance *D* and the incenter *r*_*_ (that determine the distribution of distance to the edge of coexistence) can be used to study the contributions and contextual roles of species in maintaining coexistence. To understand why, we can start with the incenter components

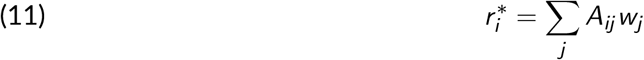

and see that it can be interpreted as a measure of the strength of competition exerted by the community on species *i*, the sum of interactions felt by that species, but where each per-capita interaction term *A*_*ij*_ is weighted by the partner’s maximal sensitivity to perturbations (the terms *w*_*j*_). Here a weak interaction with a highly sensitive species (a large *w*_*j*_) can contribute more than a weak interaction with a highly stable population (a small *w*_*j*_). If 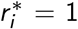, the community has a neutral effect, equal to that of the species on its own. If it is less than 1, the community facilitates that species (see supplementary figure C1). On the other hand, the distance *D* describes the edges of the feasibility domain. It reads as the inverse of an average of *S* elements (referred as *SV*_*i*_, for species vulnerability), one for each species:

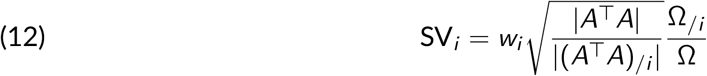

Those terms determine the distribution of coexistence states for which that species is within a certain perturbation distance from extinction. Hence, they relate to the individual vulnerability of each species.

We can combine the species-level measures 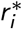 and SV_*i*_ by viewing them as the species coordinates on a two dimensional map, in other words, plotting them against each other (see Fig. 4). Intuitively, the two should be correlated: species that perceive a hostile biotic environment should also be the most vulnerable, and vice versa. This should lead to the definition of “two” particular roles: on the one hand, vulnerable and repressed, and on the other robust and facilitated. The results obtained by applying our measures to empirical data (Fig. 4) show that it doesn’t always have to be this simple, and that it is possible to define two other non-trivial qualitative “roles”.

**Figure 4.**
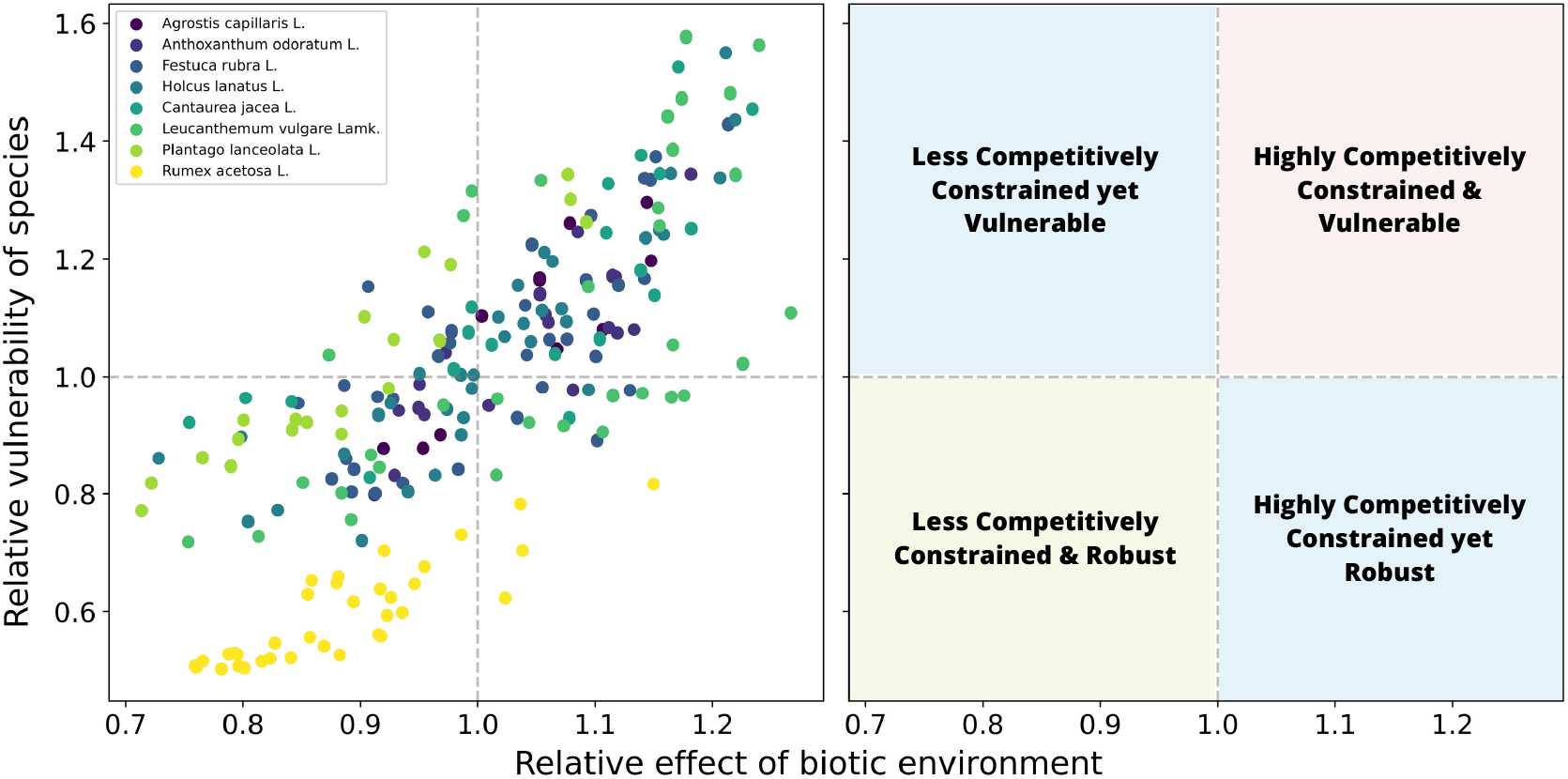
Analysis of the robustness of coexistence at the species scales. Each point on the left graph represents all the individual positions of the 8 species of the dataset within the 35 possible 4-species communities where they are present (note that in the 70 possible 4-species communities that can be generated, each species is present in 35 of them). On the x-axis, the relative effect of interactions (biotic environment) is indicated (*r*^*i*\*^ divided by the mean value for all species in the community). On the y-axis, the vulnerability of each species is indicated (SV_*i*_, divided by the average on all species of the community). This allows us to define 4 notable cases, represented on the graph on the right by the different colors.

### Application to data from a grassland experiment

Our approach characterizes the robustness of coexistence at two levels: at the scale of the community as a whole, but also at the species scale. Here, we illustrate the insights that this approach can generate for real ecological communities. We revisit data from Van Ruijven and Berendse (2009) and its subsequent analysis by Barbier et al. (2021), compiled from long-term studies of plant communities in the experimental gardens of Wageningen University, Netherlands. Here we directly use the results of Barbier et al. (2021), who estimated the interaction strengths between 8 plant species, as well as their carrying capacities. Interactions refer here to a Lotka-Volterra parametrization that differs from the one that implicitly follows from equation 1. Indeed, monocultures where used to infer species’ carrying capacities *K*_*i*_, and it is those that we consider as proxies for the abiotic conditions (and not intrinsic growth rates *r*_*i*_). The relevant interaction matrix, inferred using duo-culture experiments, follows from re-writing the L-V equations as

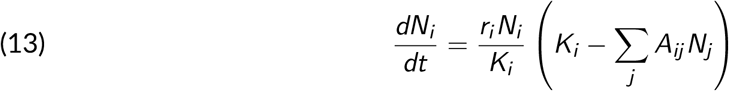

In this parametrization, *A*_*ij*_ has no dimensions and satisfies *A*_*ii*_ ≡ 1. On the basis of pairwise interaction values, we then reconstruct interaction matrices consisting of 4 species, which have been experimentally realized (Van Ruijven and Berendse, 2009). All the pairwise interaction values and carrying capacity values derived from their work are available on Zotero (See Data, script, code, and supplementary information availability section below).

To show the role of the same species in different communities, we calculated SV_*i*_ and 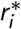 for each species within all four-species communities (See left panel of Fig. 4). We normalized these values by the mean value within each community to obtain relative species vulnerability and relative biotic effects on species, as the same species can hold different roles for the robustness of coexistence, depending on the biotic environment. Furthermore, while we unsurprisingly find the same trend as in the supplementary figure C1 (The majority of points being located in the red and green areas and being either “Highly Competitively Constrained and Vulnerable” or “Less Competitively Constrained and Robust”), we can observe non-trivial cases (blue areas of the figure). In these cases, the biotic interactions affecting the species in question are not su?cient to explain its vulnerability.

The “Less Competitively Constrained yet Vulnerable” points correspond to a case where the strong vulnerability comes from its competitive forces applied to (and not received by) other species. Indeed, to achieve coexistence, it must necessarily be of low abundance and therefore vulnerable, so that other species do not suffer too greatly from its presence. The “High Competitively Constrained yet Robust” points correspond to the case where species are useful for the coexistence of others and therefore have a high abundance (and a low vulnerability of coexistence) despite higher competitive forces experienced. These non-trivial cases explain why some points in supplementary figure C1 deviate from the expected relation.

Interestingly, the points cluster relatively well by species. This suggests that within the different 4-species communities formed by the 8 selected species, the species tend to maintain a relatively identical biotic role. Note that the abiotic environment in which these species have grown is supposedly the same. This makes ecological sense, as the biotic roles of each species depend on their phenotypic traits, and are therefore fixed by the biology of each species. For example, *Rumex Acetosa L*. is predominantly found in the green zone in Fig. 4, suggesting good persistence through low competitive forces. This fits rather well with its characterization as a weed species, present in a wide range of environments and able to coexist and persist within many ecosystems (Korpelainen and Pietiläinen, 2020).

Since the abiotic environment was assumed to be the same across the experiment, we can now determine how well or ill suited it was to particular species combinations (in terms of favouring robust coexistence). Indeed, using the carrying capacities determined during the experiments, we can determine *z*_*r*_, the minimal distance to the edge of the realized community, and *p*(*z*_*r*_), the proportion of points within this distance. This allows us to place all the communities on the *z/D*_*_ and *p*(*z*) curve (see Fig. 5). If *p*(*z*_*r*_) ≈ 1, it means that in this environment, the realized community had the most advantageous combination of biotic interactions (interaction matrix *A*) and intrinsic species parameters (carrying capacity *K*), in terms of robustness of coexistence. If *p*(*z*_*r*_), it means that this environment has led to a kind of mismatch between species interactions and species growth rates, making coexistence far less robust than what it could have been, given the set of species and their interactions.

**Figure 5.**
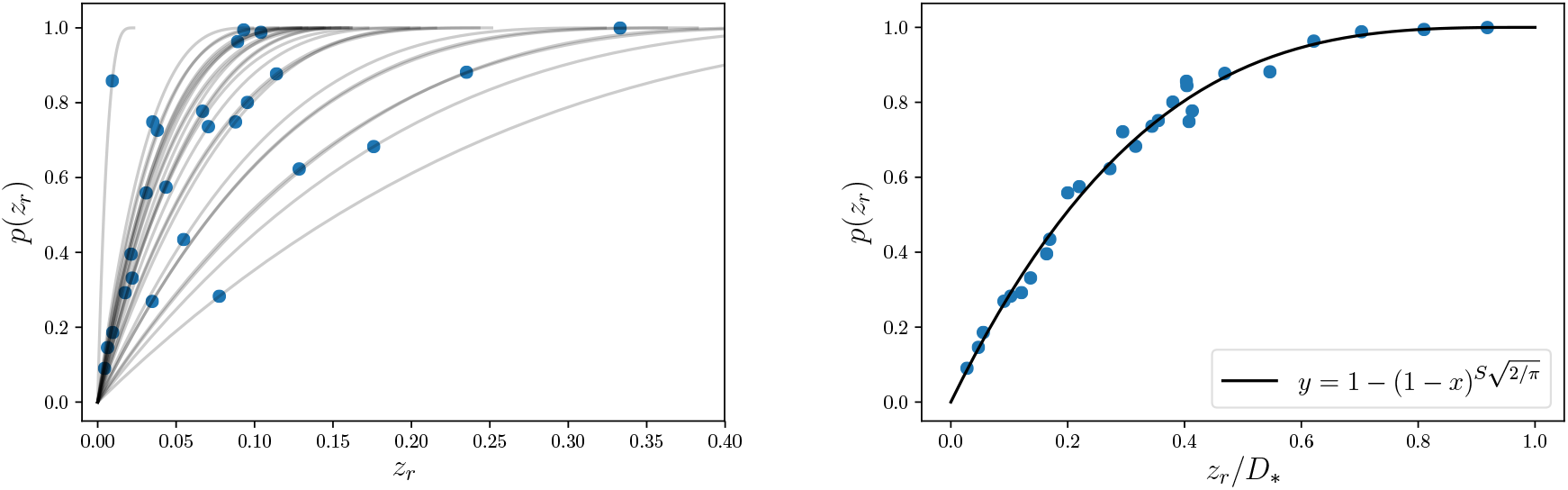
Using the empirically inferred interaction matrix between 8 plant species (Barbier et al., 2021) and their carrying capacities (taking median values for simplicity), we assembled all theoretically feasible 4-species communities (27 out of the 70 different combinations turn out to be feasible). Left: The interaction matrix for each community defines a curve, and the realized community gives the point on the curve. Large values of *z*_*r*_ (x-axis) implies high robustness (i.e. large distance from the edge of feasibility), whereas large values of *p*(*z*_*r*_) means that most communities with similar interactions are less robust. The higher this value, the better the match between the realized intrinsic parameters and biotic interactions. Right: how well suited interactions and carrying capacities go together is more clearly visualised by rescaling realized distances by the maximal distance *D*_*_. Indeed all curves collapse on a single one and we see that the communities span the whole range of *p*(*z*), meaning that some are as robust as they could be, while others are much more vulnerable than what could have been expected. The analytical graph is 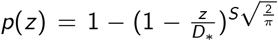 (here *S* = 4), its accuracy to predict the actual *p*(*z*) values is due to the fact that *D*_*_ ≈ *D* (see supplementary figure A2)

## Discussion

For a given community on interacting species, the function *z ↦ p* maps a value of environmental perturbation intensity *z* to the fraction *p* of coexistence states from which coexistence can be lost following such perturbations. We showed here that *p*(*z*) is a rich object to study the robustness of species coexistence, and how biotic interactions affect it, while not reducing robustness to a single number.

In Lotka-Volterra models, *p*(*z*) precisely characterizes the shape of the feasibility domain, which is the set of growth rate vectors that allow stable coexistence between all species. Indeed *p*(*z*) = ℙ (*d* ≤ *z*) determines the distribution of distances to the edge of coexistence (see Fig. 3), where for a given coexistence state, the distance to an edge corresponds to the state’s “robustness” or “full resistance”, as defined by Lepori et al. (2024) and Medeiros et al. (2021).

We showed that the function *p*(*z*) is fully parameterized by species richness 𝒮 and two characteristic distances *D* and *D*_*_, both equal to 1 in the absence of interactions. More precisely

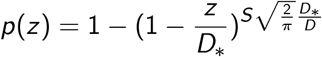

*D* is the maximal distance within the feasibility domain and thus represents the robustness of the most robust state, *r*^*^, such that *p*(*z* = *D*_*_) = 1. We derived remarkably simple formulas for *D*_*_ and *r*^*^ (See Eq. 9 and 11 and mathematical appendix), based on the interaction matrix and its inverse. Unpacking the expression for *r*^*^ allowed us to give a species-level characterization that can be interpreted as measuring the effective amount of competition that any given species feels, where its interactions are weighted by the sensitivities of its interacting partners.

The other important distance, *D*, once divided by 𝒮, determines the behaviour of *p*(*z*) at small perturbation intensity values, in the sense that 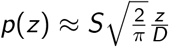 describing the edges of feasibility, which take up most of its volume if 𝒮 is large^1^. The ratio 𝒮*/D* can be used to understand how many species can be grouped together while maintaining a high percentage of robust states. More precisely, if we want to guarantee that a proportion *p* of coexistence states is robust to perturbations of intensity *ϵ*, then maximizing diversity amounts to solving

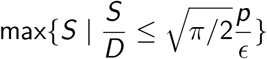

whose solution will take the form of 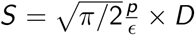, so proportional to *D*.

The expression for *D* is less simple than the one for *D*_*_, but can also be used to give complementary species-level characterization of coexistence. In line with Allen-Perkins et al. (2023), we can decompose *D* to measure the robustness of each species persistence conditioned on overall coexistence. This interpretation, together with the one relating *r*^*^ to effective competition pressure, can be used to reveal the contextual roles of species in maintaining coexistence. The biotic context created within a coexisting community can be favorable or unfavorable to individual species through the balance of interactions they receive and emit and how hostile they are to others (See different panels of Fig. 4). It is interesting to note that the species present in the dataset used in the study seem to retain relatively the same role regardless of community composition. It would be interesting to extend this analysis to larger datasets to study the consistency of species roles in maintaining robust coexistence. If we consider the contribution to the community-scale robustness of coexistence as a function rendered by a species within the community, it is likely that certain species correspond to “key species” (Power et al., 1996; Whittaker and Cottee-Jones, 2012).

Broadly speaking, our theory highlights a negative effect, amplified by species richness, of the intensity of the interaction forces and the sensitivity of the species on the robustness of coexistence. Figure 4 and supplementary figure C1 also show the relationship between strong inter-specific competition faced by species and their vulnerability of coexistence. These results are consistent with the existing literature on the effect of interactions on community coexistence or stability under environmental perturbations (Barabás et al., 2016; Chesson, 2000; Hale et al., 2020; Mccann et al., 1998; Vallina and Le Quéré, 2011). The fact that features of the inverse interaction matrix are present in both *D* and *D*_*_ highlights the importance of network structure, as the inverse matrix encodes net effects between species, via all indirect interaction pathways. For the same overall mean interaction strength, net effects can be very different depending on the way the matrix *A* is organized. This is consistent with previous research on the effect of network structure on coexistence (especially in cases with more than two species) as on other stability notions (Barabás et al., 2016; Cenci et al., 2018b; Lurgi et al., 2016; Serván et al., 2018). This leads to an important ecological conclusion: vulnerability to extinction depends on how a species is affected by others through direct interactions, combined with the sensitivities of those species (how they amplify environmental change). Here sensitivity is a potentially collective notion that arises from indirect interactions between species, and is thus sensitive to the interaction structure.

As in previous studies of asymmetry of the feasibility domain, our theory strongly depends on the way environmental disturbances are modeled (Allen-Perkins et al., 2023; Cenci et al., 2018a; De Laender et al., 2023; Lepori et al., 2024). This highlights the importance of taking into account the type of disturbance when studying the stability of a community (Arnoldi et al., 2018; Arnoldi et al., 2019; Bender et al., 1984) and suggests that different results could be obtained by considering other types of disturbance (ie. that vary through time, and/or scale with species standing biomass). Deepening our theory to account for more general types of disturbance could be an interesting direction.

Coexistence is defined as the maintenance of positive abundance of all species in a community. No attention is paid to total biomass, ecosystem functions, turnover, or processes at the meta-community level. Our results should therefore not be interpreted as evidence of a negative effect of biodiversity on stability in the sense of maintaining biomass or ecological function over time (Loreau and Mazancourt, 2013), nor on the resistance or resilience of the the community (Arnoldi et al., 2016; Kéfi et al., 2019). It simply highlights the di?culty for complex interaction networks to generate communities that can tolerate environmental disturbances without losing any species. This vision of a fixed community and coexistence seen as the absolute persistence of all species over time is, however, clearly limited and open to criticism. It would be interesting to develop approaches that include turnover or variations in species interactions over time.

Another caveat is the supposed independence between biotic and abiotic parameters. This unrealistic assumption means that a change in abiotic environmental conditions (disruption of growth rates or carrying capacity) should not change biotic interactions. This assumption is necessary to define the feasibility domain (Saavedra et al., 2017). However, the empirical applications we present (determination of the biotic role of different species within several communities; quantification of the adequation between a given abiotic environment and a certain biotic assemblage) illustrate how to overcome this issue. Indeed, in the experimental data, the abiotic environment is the same for each community studied and is not subject to change.

Overall, this study provides an understanding of the link between the conditions under which communities coexist and the robustness of this coexistence. On the one hand, the analytical results provide a clear explanation of the relationships between the various mathematical elements involved in feasibility domain analysis. On the other hand, they enable us to link the interpretations made specifically through the analysis of the notion of feasibility domain to more general notions of community ecology. In doing so, we have linked different measures of stability and placed the robustness of coexistence within the multidimensional concept of ecological stability (Donohue et al., 2016; Radchuk et al., 2019).

## Supporting information

Mathematical Appendix

## Acknowledgements

We thank Matthieu Barbier for the data provided and useful discussions. We also thank Rudolf Rohr and Nicolas Loeuille for inspiring discussions throughout the article writing process. We finally thanks reviewers and editor for their very constructive feedback, valuable suggestions and timely handling of our manuscript. This preprint version of this article has been peer-reviewed and recommended by Peer Community In Ecology (https://doi.org/10.24072/pci. ecology.100681; De Laender, 2024).

## Data, script, code, and supplementary information availability

Script and data useful for “Application to data from a grassland experiment” section are available online: 10.5281/zenodo.10534234; For more information on the dataset, please contact Barbier et al. (2021); Desallais et al., 2024b

Supplementary information, including appendices A, B and C and a mathematical appendix, is available online : (https://zenodo.org/doi/10.5281/zenodo.12744286; Desallais et al., 2024a

## Footnotes

1 This last remark is only a geometrical way of saying that for many interacting species, in the absence of prior knowledge of abiotic conditions, there is a high chance that at least one of those species is close to extinction.

## References

Abrams (1984). Variability in resource consumption rates and the coexistence of competing species. Theoretical Population Biology 25, 106–124. 10.1016/0040-5809(84)90008-X. URL: https://www.sciencedirect.com/science/article/pii/004058098490008X.

Abrams Brassil, Holt (2003). Dynamics and responses to mortality rates of competing predators undergoing predator–prey cycles. Theoretical Population Biology 64, 163–176. 10.1016/S0040-5809(03)00067-4. URL: https://linkinghub.elsevier.com/retrieve/pii/S0040580903000674 (visited on 01/09/2023).

Allen-Perkins A, García-Callejas D, Bartomeus I, Godoy O (2023). Structural asymmetry in biotic interactions as a tool to understand and predict ecological persistence. Ecology Letters n/a. 10.1111/ele.14291.

Armstrong RA, McGehee R (1976). Coexistence of species competing for shared resources. Theoretical Population Biology 9, 317–328. 10.1016/0040-5809(76)90051-4. URL: https://linkinghub.elsevier.com/retrieve/pii/0040580976900514 (visited on 02/08/2022).

Arnoldi JF, Loreau M, Haegeman B (2016). Resilience, reactivity and variability: A mathematical comparison of ecological stability measures. Journal of Theoretical Biology 389, 47–59. 10.1016/j.jtbi.2015.10.012. URL: https://linkinghub.elsevier.com/retrieve/pii/S0022519315005056 (visited on 02/08/2022).

Arnoldi JF, Bideault A, Loreau M, Haegeman B (2018). How ecosystems recover from pulse perturbations: A theory of short-to long-term responses. Journal of theoretical biology 436, 79–92. 10.1016/j.jtbi.2017.10.003. URL: https://www.ncbi.nlm.nih.gov/pmc/articles/PMC5675055/ (visited on 01/08/2023).

Arnoldi JF, Loreau M, Haegeman B (2019). The inherent multidimensionality of temporal variability: how common and rare species shape stability patterns. Ecology letters 22, 1557–1567. 10.1111/ele.13345. URL: https://www.ncbi.nlm.nih.gov/pmc/articles/PMC6756922/ (visited on 07/18/2022).

Barabás G, J. Michalska-Smith M, Allesina S (2016). The Effect of Intra-and Interspecific Competition on Coexistence in Multispecies Communities. The American Naturalist 188. Publisher: The University of Chicago Press, E1–E12. 10.1086/686901. URL: https://www.journals.uchicago.edu/doi/full/10.1086/686901 (visited on 07/25/2022).

Barbier M, Mazancourt C, Loreau M, Bunin G (2021). Fingerprints of High-Dimensional Coexistence in Complex Ecosystems. Physical Review X 11. Publisher: American Physical Society, 011009. 10.1103/PhysRevX.11.011009. URL: https://link.aps.org/doi/10.1103/PhysRevX.11.011009 (visited on 07/18/2022).

Bartomeus I, Saavedra S, Rohr RP, Godoy O (2021). Experimental evidence of the importance of multitrophic structure for species persistence. Proceedings of the National Academy of Sciences 118, e2023872118. 10.1073/pnas.2023872118. URL: http://www.pnas.org/lookup/doi/10.1073/pnas.2023872118 (visited on 02/08/2022).

Bender EA, Case TJ, Gilpin ME (1984). Perturbation Experiments in Community Ecology: Theory and Practice. Ecology 65, 1–13. 10.2307/1939452. URL: http://doi.wiley.com/10.2307/1939452 (visited on 02/08/2022).

Brose U, Williams RJ, Martinez ND (2006). Allometric scaling enhances stability in complex food webs. Ecology Letters 9, 1228–1236. 10.1111/j.1461-0248.2006.00978.x.

Cenci S, Montero-Castaño A, Saavedra S (2018a). Estimating the effect of the reorganization of interactions on the adaptability of species to changing environments. Journal of Theoretical Biology 437, 115–125. 10.1016/j.jtbi.2017.10.016. URL: https://www.sciencedirect.com/science/article/pii/S0022519317304794.

Cenci S, Song C, Saavedra S (2018b). Rethinking the importance of the structure of ecological networks under an environment-dependent framework. Ecology and Evolution 8, 6852–6859. 10.1002/ece3.4252. URL: https://onlinelibrary.wiley.com/doi/abs/10.1002/ece3.4252 (visited on 07/25/2022).

Chesson P (2000). Mechanisms of Maintenance of Species Diversity. Annual Review of Ecology and Systematics 31, 343–366. 10.1146/annurev.ecolsys.31.1.343. URL: https://www.annualreviews.org/doi/10.1146/annurev.ecolsys.31.1.343 (visited on 02/08/2022).

Coulson T, Kendall BE, Barthold J, Plard F, Schindler S, Ozgul A, Gaillard JM (2017). Modeling Adaptive and Nonadaptive Responses of Populations to Environmental Change. The American Naturalist 190. PMID: 28829647, 313–336. 10.1086/692542. eprint: https://doi.org/10.1086/692542. URL: https://doi.org/10.1086/692542.

De Laender F (2024). How environmental perturbations affect coexistence. 10.24072/pci.ecology.100681. URL: https://doi.org/10.24072/pci.ecology.100681.

De Laender F, Carpentier C, Carletti T, Song C, Rumschlag SL, Mahon MB, Simonin M, Meszéna G, Barabás G (2023). Mean species responses predict effects of environmental change on coexistence. Ecology Letters 26, 1535–1547. 10.1111/ele.14278. eprint: https://onlinelibrary.wiley.com/doi/pdf/10.1111/ele.14278. URL: https://onlinelibrary.wiley.com/doi/abs/10.1111/ele.14278.

Deng J, Taylor W, Saavedra S (2022). Understanding the impact of third-party species on pairwise coexistence. PLOS Computational Biology 18, 1–21. 10.1371/journal.pcbi.1010630. URL: https://doi.org/10.1371/journal.pcbi.1010630.

Desallais M, Loreau M, Arnoldi JF (2024a). Appendices of the “The distribution of distances to the edge of species coexistence” scientific article. 10.5281/zenodo.12744286. URL: https://doi.org/10.5281/zenodo.12744287.

Desallais M, Loreau M, Jean-François A (2024b). Functions and scripts used on the “The distribution of distances to the edge of species coexistence” scientific article. 10.5281/zenodo.10534234. URL: https://doi.org/10.5281/zenodo.10905742.

Donohue I, Hillebrand H, Montoya JM, Petchey OL, Pimm SL, Fowler MS, Healy K, Jackson AL, Lurgi M, McClean D, O’Connor NE, O’Gorman EJ, Yang Q (2016). Navigating the complexity of ecological stability. Ecology Letters 19. Ed. by Frederick Adler, 1172–1185. 10.1111/ele.12648. URL: https://onlinelibrary.wiley.com/doi/10.1111/ele.12648 (visited on 02/08/2022).

Grilli J, Adorisio M, Suweis S, Barabás G, Banavar JR, Allesina S, Maritan A (2017). Feasibility and coexistence of large ecological communities. Nature Communications 8, 14389. 10.1038/ncomms14389. URL: http://www.nature.com/articles/ncomms14389 (visited on 02/08/2022).

Hale KRS, Valdovinos FS, Martinez ND (2020). Mutualism increases diversity, stability, and function of multiplex networks that integrate pollinators into food webs. Nature Communications 11, 2182. 10.1038/s41467-020-15688-w. URL: https://doi.org/10.1038/s41467-020-15688-w.

Hastings A (1980). Disturbance, coexistence, history, and competition for space. Theoretical Population Biology 18, 363–373. 10.1016/0040-5809(80)90059-3. URL: https://linkinghub.elsevier.com/retrieve/pii/0040580980900593 (visited on 02/08/2022).

Hutchinson GE (1961). The Paradox of the Plankton. THE AMERICAN NATURALIST, 9.

Kéfi S, Domínguez-García V, Donohue I, Fontaine C, Thébault E, Dakos V (2019). Advancing our understanding of ecological stability. Ecology Letters 22. Ed. by Tim Coulson, 1349–1356. 10.1111/ele.13340. URL: https://onlinelibrary.wiley.com/doi/10.1111/ele.13340 (visited on 02/07/2022).

Korpelainen H, Pietiläinen M (2020). Sorrel (Rumex acetosa L.): Not Only a Weed but a Promising Vegetable and Medicinal Plant. The Botanical Review 86, 234–246. 10.1007/s12229-020-09225-z. URL: https://doi.org/10.1007/s12229-020-09225-z.

Lepori VJ, Loeuille N, Rohr RP (2024). Robustness versus productivity during evolutionary community assembly: short-term synergies and long-term trade-offs. Proceedings of the Royal Society B: Biological Sciences 291, 20232495. 10.1098/rspb.2023.2495. eprint: https://royalsocietypublishing.org/doi/pdf/10.1098/rspb.2023.2495. URL: https://royalsocietypublishing.org/doi/abs/10.1098/rspb.2023.2495.

Levins R (1968). Evolution in Changing Environments: Some Theoretical Explorations. Princeton University Press. URL: https://books.google.fr/booksfiid=yOQ9DwAAQBAJ&lr=&source=gbs_navlinks_s.

Loreau M, Mazancourt C (2013). Biodiversity and ecosystem stability: a synthesis of underlying mechanisms. Ecology Letters 16. Ed. by Emmett Duffy, 106–115. 10.1111/ele.12073. URL: https://onlinelibrary.wiley.com/doi/10.1111/ele.12073 (visited on 02/07/2022).

Lurgi M, Montoya D, Montoya JM (2016). The effects of space and diversity of interaction types on the stability of complex ecological networks. Theoretical Ecology 9, 3–13. 10.1007/s12080-015-0264-x. URL: https://doi.org/10.1007/s12080-015-0264-x (visited on 08/12/2022).

Mccann K, Hastings A, Huxel G (1998). Weak Trophic Interactions and the Balance of Nature. Nature 395, 794–798. 10.1038/27427.

Medeiros LP, Song C, Saavedra S (2021). Merging dynamical and structural indicators to measure resilience in multispecies systems. Journal of Animal Ecology 90, 2027–2040. 10.1111/1365-2656.13421. URL: https://onlinelibrary.wiley.com/doi/10.1111/1365-2656.13421 (visited on 02/08/2022).

Meszéna G, Gyllenberg M, Pásztor L, Metz JA (2006). Competitive exclusion and limiting similarity: A unified theory. Theoretical Population Biology 69, 68–87. 10.1016/j.tpb.2005.07.001. URL: https://www.sciencedirect.com/science/article/pii/S004058090500095X.

Otto SB, Rall BC, Brose U (2007). Allometric degree distributions facilitate food-web stability. Nature 450, 1226–1229. 10.1038/nature06359.

Power ME, Tilman D, Estes JA, Menge BA, Bond WJ, Mills LS, Daily G, Castilla JC, Lubchenco J, Paine RT (1996). Challenges in the Quest for Keystones: Identifying keystone species is dificult—but essential to understanding how loss of species will affect ecosystems. BioScience 46, 609–620. 10.2307/1312990. eprint: https://academic.oup.com/bioscience/article-pdf/46/8/609/650270/46-8-609.pdf. URL: https://doi.org/10.2307/1312990.

Radchuk V, Laender FD, Cabral JS, Boulangeat I, Crawford M, Bohn F, Raedt JD, Scherer C, Svenning JC, Thonicke K, Schurr FM, Grimm V, Kramer-Schadt S (2019). The dimensionality of stability depends on disturbance type. Ecology Letters 22. Ed. by Ian Donohue, 674–684. 10.1111/ele.13226. URL: https://onlinelibrary.wiley.com/doi/10.1111/ele.13226 (visited on 02/01/2022).

Ribando JM (2006). Measuring Solid Angles Beyond Dimension Three. Discrete & Computational Geometry 36, 479–487. 10.1007/s00454-006-1253-4. URL: http://link.springer.com/10.1007/s00454-006-1253-4 (visited on 02/08/2022).

Rohr RP, Saavedra S, Bascompte J (2014). On the structural stability of mutualistic systems. Science 345, 1253497. 10.1126/science.1253497. URL: https://www.science.org/doi/10.1126/science.1253497 (visited on 02/08/2022).

Rohr RP, Saavedra S, Peralta G, Frost CM, Bersier LF, Bascompte J, Tylianakis JM (2016). Persist or Produce: A Community Trade-Off Tuned by Species Evenness. The American Naturalist 188. PMID: 27622875, 411–422. 10.1086/688046. eprint: https://doi.org/10.1086/688046. URL: https://doi.org/10.1086/688046.

Roughgarden J (1975). A Simple Model for Population Dynamics in Stochastic Environments. The American Naturalist 109, 713–736. URL: http://www.jstor.org/stable/2459866.

Saavedra S, Rohr RP, Bascompte J, Godoy O, Kraft NJB, Levine JM (2017). A structural approach for understanding multispecies coexistence. Ecological Monographs 87, 470–486. 10.1002/ecm.1263. URL: https://onlinelibrary.wiley.com/doi/10.1002/ecm.1263 (visited on 02/07/2022).

Serván CA, Capitán JA, Grilli J, Morrison KE, Allesina S (2018). Coexistence of many species in random ecosystems. Nature Ecology & Evolution 2. Number: 8 Publisher: Nature Publishing Group, 1237–1242. 10.1038/s41559-018-0603-6. URL: https://www.nature.com/articles/s41559-018-0603-6 (visited on 07/25/2022).

Song C, Altermatt F, Pearse I, Saavedra S (2018a). Structural changes within trophic levels are constrained by within-family assembly rules at lower trophic levels. Ecology Letters 21. Ed. by Josè Marìa Gomez, 1221–1228. 10.1111/ele.13091. URL: https://onlinelibrary.wiley.com/doi/10.1111/ele.13091 (visited on 02/08/2022).

Song C, Rohr RP, Saavedra S (2018b). A guideline to study the feasibility domain of multi-trophic and changing ecological communities. Journal of Theoretical Biology 450, 30–36. 10.1016/j.jtbi.2018.04.030. URL: https://linkinghub.elsevier.com/retrieve/pii/S0022519318302042 (visited on 02/08/2022).

Vallina SM, L. Quéré C (2011). Stability of complex food webs: Resilience, resistance and the average interaction strength. Journal of Theoretical Biology 272, 160–173. 10.1016/j.jtbi.2010.11.043. URL: https://www.sciencedirect.com/science/article/pii/S0022519310006387 (visited on 07/25/2022).

Van Ruijven J, Berendse F (2009). Long-term persistence of a positive plant diversity–productivity relationship in the absence of legumes. Oikos 118, 101–106. 10.1111/j.1600-0706.2008.17119.x. URL: https://onlinelibrary.wiley.com/doi/abs/10.1111/j.1600-0706.2008.17119.x (visited on 12/23/2023).

Volterra V (1926). Fluctuations in the Abundance of a Species considered Mathematically1. Nature 118, 558–560. 10.1038/118558a0. URL: https://www.nature.com/articles/118558a0 (visited on 07/15/2022).

Whittaker R, Cottee-Jones HEW (2012). The keystone species concept: a critical appraisal. Frontiers of Biogeography 4, 117–127. 10.21425/F54312533.

Williams RJ (2008). effects of network and dynamical model structure on species persistence in large model food webs. Theoretical Ecology 1, 141–151. 10.1007/s12080-008-0013-5. URL: http://link.springer.com/10.1007/s12080-008-0013-5 (visited on 07/19/2022).

