## Appendix A for "The distribution of distances to the edge of species coexistence"

### Appendix A
### The distribution of distances to the edge of species coexistence
### Relation between $\Omega$, $D$ and $D_*$

Mario Desallais, Michel Loreau, and Jean-François Arnoldi

In this study, we highlight the importance of taking into account the shape of the feasibility domain and its size to characterize the robustness of coexistence induced by species interactions. Two different measures, therefore, emerge: $\Omega$, a proxy for the probability of coexistence, and $D$ (or $D_*$), a proxy for the robustness of coexistence. However, these are not independent. Fig. 1 shows the relationship between the two values.

**Figure 1** – Relation between $D$ and $\Omega^{2/S}$ for random matrices of size $S$ x $S$. If $D$ controls the distribution of distance to the edge of feasibility, $\Omega$ corresponds to a relative volume of the feasibility domain. The two notions are not equivalent, but still are closely related, which is seen here by comparing D to $\Omega^{2/S}$.

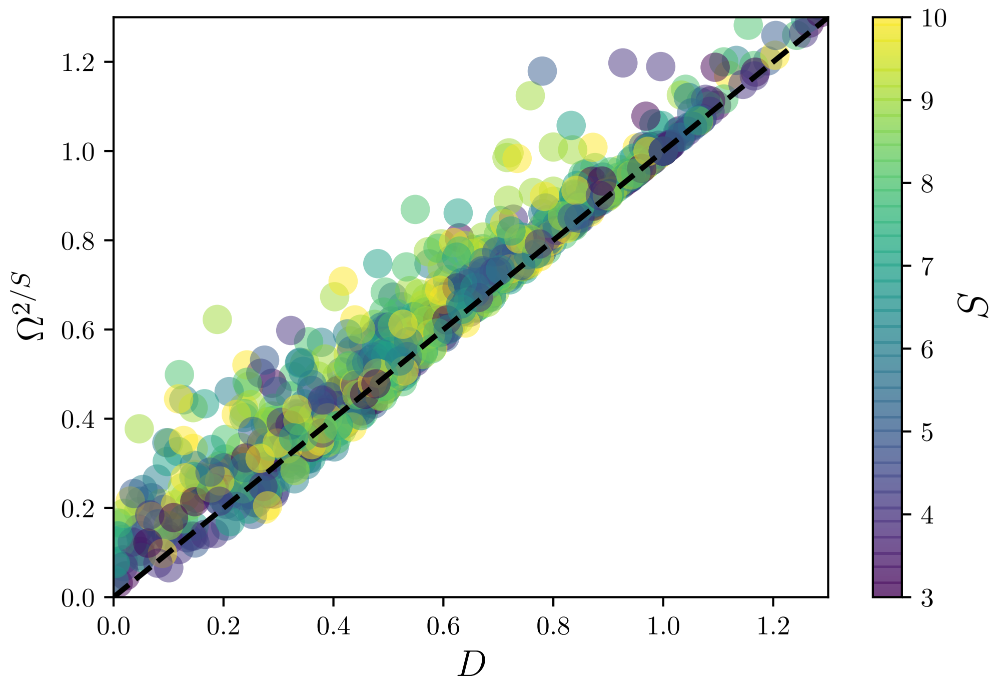


Interestingly, $D$ and $D_*$ are also very closely related. This is somewhat visible in their respective expressions, and confirmed numerically (see Fig. 2). This a useful thing to note because $D_*$ is much simpler to compute, interpret, and manipulate than $D$, although it is the latter that is expected to driver of the major part of the function $p(\hat{z})$, at least when considering species-rich communities.

#### References

Barbier M, Mazancourt C, Loreau M, Bunin G (2021). *Fingerprints of High-Dimensional Coexistence in Complex Ecosystems*. *Physical Review X* **11**. Publisher: American Physical Society, 011009. https://doi.org/10.1103/PhysRevX.11.011009. URL: https://link.aps.org/doi/10.1103/PhysRevX.11.011009 (visited on 07/18/2022).

**Figure 2** – Correlation between robustness of coexistence $D$ and characteristic distance $D_*$. Left panel shows this correlation for randomly generated matrices of variable size ($S$ between 3 and 11). The right panel shows this correlation for matrices from real communities ($S$=4), based on the Barbier et al. (2021) dataset (see section "Application to data from grassland experiment" from the main article). The diagonal blue line corresponds to the x=y line in both cases. Note that D and D* are indeed closely related, for random matrices as for empirical matrices.

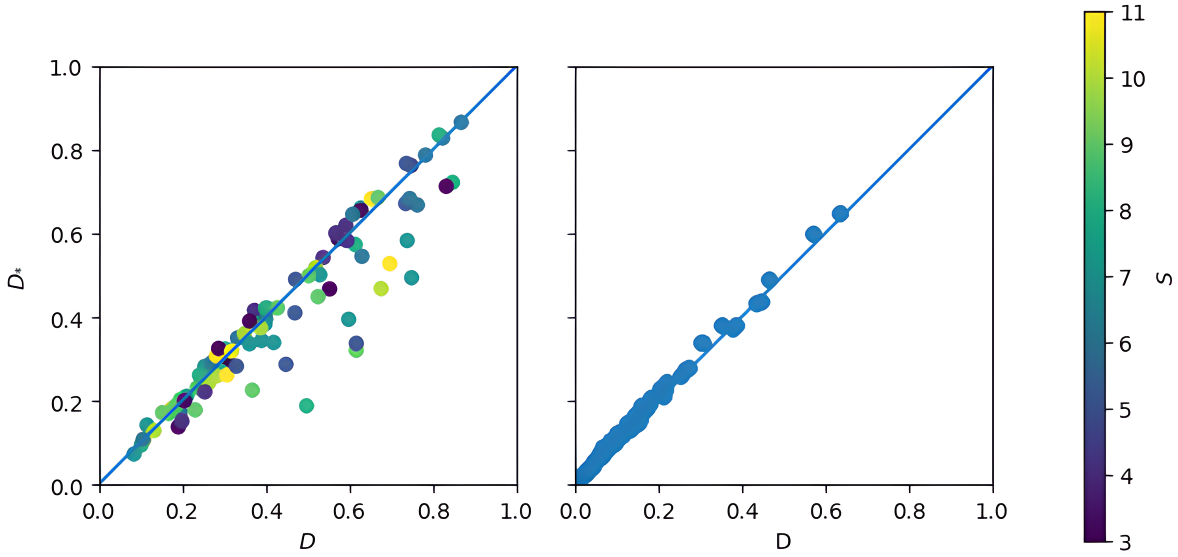
