## Appendix B for "The distribution of distances to the edge of species coexistence"

### Appendix B
### The distribution of distances to the edge of species coexistence
### Persistence of species in simulated ecological systems

Mario Desallais, Michel Loreau, and Jean-François Arnoldi

In this appendix, we investigate the relationship between the metrics proposed in this study (See Eq. 10 and 11 of the main article) and actual species and community persistence. Although our main results consist in the distribution function of distances to the edge of species coexistence and the characterization of biotic roles at the species level in this coexistence, this verification remains essential to ensure the relevance of speaking about persistence, robustness or vulnerability of coexistence.

In the absence of an adequate dataset for this test, we employ numerical simulations using a generalized Lotka-Volterra model (See Eq. 1 of the main article) with random parameters. The simulation we used for this test follows this procedure : We generate $n$ interaction matrices $A$ of size $S \times S$, ensuring the are D-stable. We then sample a single set of growth rates (vector $r$). By construction, this defines $n$ sets of species abundances (recall, $N^*(r) = A^{-1}r$) and therefore $n$ communities. The coexistence and feasability of the latter depend on wheter the vector $r$ is within the feasibilty domain defined by the interaction matrix $A$ associated.

To simulate a changing environment, we randomly perturb the growth rate vector $r$, applying a small $\delta r_i$ at each time step of the simulation. Consequently, at each time step, new species abundances $N_i$ are defined. Coexistence occurs if $r$ lies within the feasibility domain and all $N_i > 0$, otherwise it does not. When a community coexists for at least one time step, this defines a stable coexistence period. This period ends if, at any subsequent time step, any abundance falls to zero or below.

At the end of the simulation, we calculate the average duration of these "stable coexistence periods" for each community, over the total duration of the simulation. These values are represented on the Y axis of the left-hand panel in Figure 1 and we hypothesize that $D_*$ is a good predictor of these values.

During the simulation, we also count each coexistence loss event for each community. Since each coexistence loss can be associated with a species (the one whose abundance falls to 0 or below), we calculate the proportion of extinction / coexistence loss events associated with each species in the community. This defines the Y axis of the right-hand panel in figure 1. We hypothesize that the value of $SV_i$ for each species (relative to the mean in the community) predicts this proportion.

These two hypotheses are verified, the results of which are shown in Figure 1. A large value of $D_*$ can allow a relatively long average duration of stable coexistence (although this does not guarantee it), while a small value of $D_*$ forces a small average duration of coexistence. Regarding species extinctions, a low value of $SV_i$ seems to guarantee a lower risk of extinction throughout the simulation, while a high value of $SV_i$ are associated with a high proportion of extinction. This is particularly true for extreme values. The results obtained for $SV_i$ values (relative to the community average) close to 1 are expected given the randomness of disturbances on $\delta_r$ and the fact that at $SV_i$ close to 1, the species concerned is neither particularly favoured nor particularly damaged by other species.

The results obtained (1) are particularly sensitive to the parameterization of the simulation (especially, total time of the random walk, initial value of $r_i$, and value taken by the $\delta r_i$ during the random walk), but the conclusions drawn above are consistent over the simulations. Nonetheless, these simulations should only be considered as a first check, and deserve to be further explored to provide more solid proof of the link between distances to the edges of the feasibility domain and persistence. In this sense, it would be very interesting to extend this study with a dataset from real experiment that quantified biotic interactions between species and the persistence of communities and species in the face of disturbance.

**Figure 1** – Simulations-based testing of the link between community persistence (left panel) or species persistence (right panel) and measures of coexistence derived from the $p(z)$ function. Y-axis of left panel is displayed in log scale. Here, $n = 1000$, $S = 3$, non-diagonal elements of $A$ are comprise between $-1$ and $1$ and $\delta r_i$ is drawn in a normal distribution with mean $0$ and variance $1$.

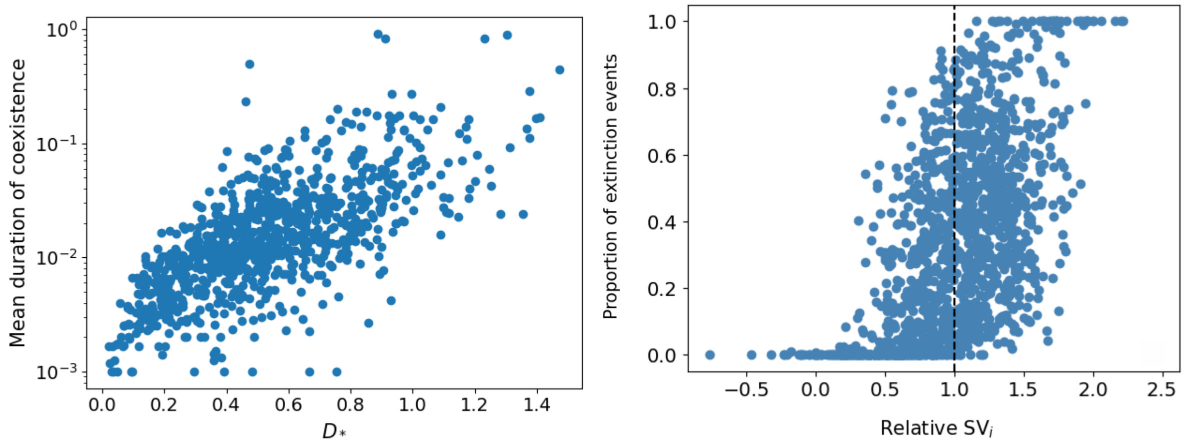
