## Appendix C for "The distribution of distances to the edge of species coexistence"

### Appendix C
#### The distribution of distances to the edge of species coexistence
#### Role of absolute interaction strength in species vulnerability

**Mario Desallais, Michel Loreau, and Jean-François Arnoldi**

Although in the main article we used the $SV_i$ and $r_i^*/D_*$ measures relative to their average in the community, it is also possible to use and compare them in absolute terms. By doing this, a strong correlation between these two values can be observed (Fig. 1). This suggests that species that are constrained by others (through competition, highlighted in red in Fig. 1) are generally the ones that are mostly vulnerable. Conversely, species that tend to benefit from others (through facilitation, highlighted in green in Fig. 1) are those that are less vunerable.

**Figure 1** – Correlation between the vulnerability of each species within a community ($SV_i$) and the effect of the biotic environment (interaction between species) on each ($r_i^*/D_*$). Each point represents one species within a community of 10 species (500 points in total). The vertical dot line corresponds to x=1, the qualitative threshold of the biotic effect on species. If this value is less than 1 (green box on the figure), this implies that the biotic environment is overall favorable (facilitating) to the concerned species. If upper than 1 (red box on the figure), it implies that the biotic environment is overall unfavorable through competition subjected to the species. Spearman rank correlation = 0.67 ; associated p-value : $2.5e^{-68}$

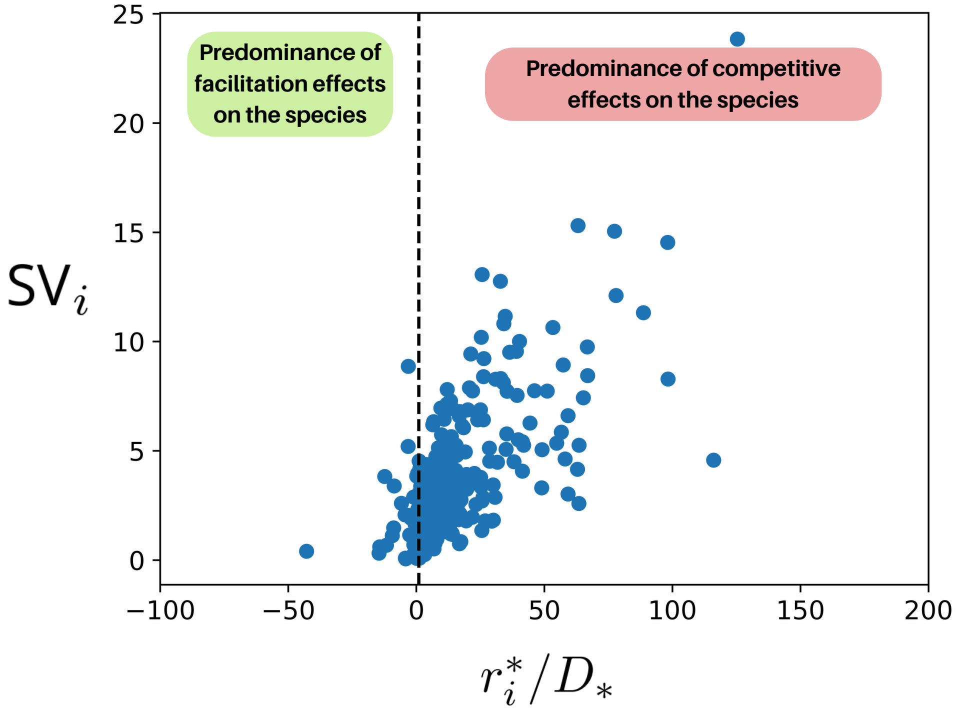
