## Supplementary material for "The distribution of distances to the edge of species coexistence": Mathematical Appendix

### Supplementary material
### The distribution of distances to the edge of species coexistence

Mario Desallais, Michel Loreau, and Jean-Francois Arnoldi

#### 1 Established theory

Consider a Generalized Lotka-Volterra model written as

$$\frac{1}{N_i}\frac{dN_i}{dt} = r_i - \sum_{j=1}^{S} A_{ij}N_j; A_{ii} > 0; i = 1, \dots, S$$

A growth rate vector $r$ is feasible if the fixed point $N(r) = A^{-1}r$ is interior (all its components are positive). We do not worry about the stability of this state -additional conditions on the interaction matrix $A$ (D-stability) can guarantee that any interior fixed point is automatically stable. Multiplying the vector $r$ by some positive constant does not change feasibility: the latter is determined by the direction of $r$, not by its magnitude. The feasibility domain $D_f(A)$ is the subspace of all growth rate directions (a hyper-sphere) that are feasible, it is a finite geometrical object more precisely a convex polytope (but we will come back to this). Remarkably, we can easily compute the relative volume of this subspace, which defines a probability measure $\mathbb{P}(D_f) \leq 1$. For that, we note that drawing growth rate values $r_i$ from independent standard normal distributions defines a uniform sampling of the directions of growth rate vectors. Integrating over all feasible configurations amounts to computing the gaussian integral and, after a change of variables, gives

$$\mathbb{P}(D_f) = \frac{1}{\sqrt{2\pi}^S}\int_{N(r)\in\mathbb{R}_+^S} e^{-\frac{\|r\|^2}{2}} d^S r = \frac{|A|}{\sqrt{2\pi}^S}\int_{\mathbb{R}_+^S} e^{-\frac{\|AN\|^2}{2}} d^S N$$

where we changed variables to integrate in $N$-space via $N = A^{-1}r$. Thus $\mathbb{P}(D_f)$ is simply the cumulative distribution noted $\Phi_{A^\top A}(0)$ of a multivariate normal distribution centered on 0 and with covariance matrix $C = \left(A^\top A\right)^{-1}$. In the absence of interactions $\mathbb{P}(D_f) = 2^{-S}$ so to focus on the effect of interactions it is convenient to define a ratio of probabilities, namely

$$\Omega\left(A^\top A\right) = \frac{\mathbb{P}(D_f(A))}{2^{-S}} = 2^S \times \Phi_{A^\top A}(0)$$

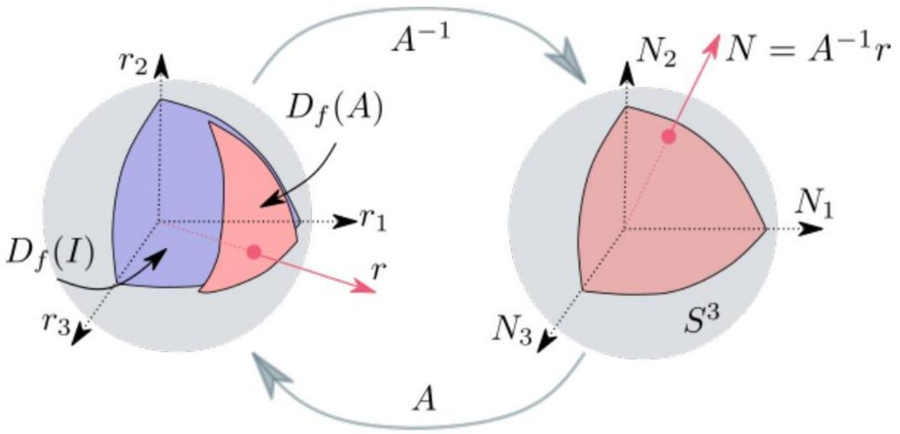


Figure 1: The feasibility domain $D_f(A)$ (in light red on the left) is defined as the subset of growth rate directions that, given a pair-wise interaction matrix $A$, allows coexistence between all species. It is the intersection of the sphere with the image in $r$-space (via the matrix $A$ ) of the positive quadrant in $N$-space (shown on the right). In the absence of interactions, the feasibility domain is the intersection of the positive quadrant and the sphere (in blue). The probability of feasibility $\mathbb{P}(D_f)$ is the ratio between the volume of $D_f$ and the volume of that sphere.

As previously mentioned, $D_f$ is a convex polytope: the generalization of a triangle (with $S$ vertices instead of 3 , connected by $S(S-1)/2$ edges and drawn on the sphere instead of the plane). This can be understood by realizing that the i-th column of $A$ defines the growth rate vector $r^{(i)}$ such that species $i$ has unit abundance while all others are exactly at 0 . Such points, when represented on the sphere are the $S$ vertices of $D_f(A)$ and any feasible growth rate can be written as a positive linear combination of the extreme vectors $r^{(i)}$. In particular, the path $\phi r^{(i)} + (1-\phi)r^{(j)}$, where $0 \leq \phi \leq 1$, and $i \neq j$, once projected on the sphere draws an edge of the domain (only species $i$ and $j$ have non zero abundance).

These first results are nothing new, but will serve as an introduction to our subsequent analysis, where we want to go beyond the volume of $D_f$ but also describe relevant features of its shape.

#### 2 Motivation

We are interested in the fragility of coexistence, stated in the following way: If a growth rate vector is feasible, how easy is it to 'push it' out of the feasibility domain? This depends on how close to the boundary that vector was in the first

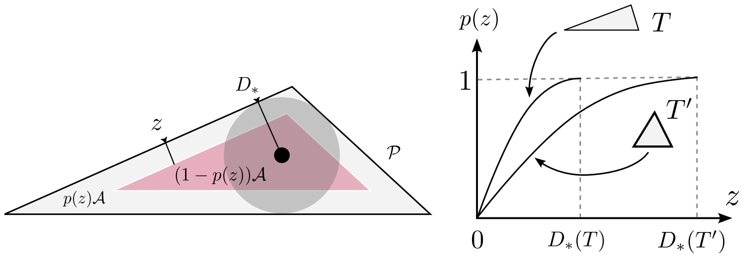


Figure 2: Left: Triangles are parametrized by their area $\mathcal{A}$ and perimeter $\mathcal{P}$. $D_*$ is the radius of the largest circle that can be contained in the triangle. For a given value $z \leq D_*$, the fraction $1 - p(z)$ of points that lie further than a euclidian distance $z$ from an edge is the relative area of an inscribed triangle, whose own edges are precisely at distance $z$ from those of the original triangle. Right: the fraction $p(z)$ entirely describes the distribution of distances since $p(z) = \mathbb{P}(\text{ distance } \leq z)$, and is fully determined by the value $z/D_*$. For a fixed area the optimal shape is those of equilateral triangles which maximise $D_*$.

place. To deduce a general statement about the interaction matrix we should instead ask: how likely is it that points in $D_f(A)$ are closer to the boundary than some threshold value $z$ ? What we will do is determine a characteristic distance $D_*$ such that, given a prescribed distance $z, z/D_*$ determines the proportion of points that lie at that distance from the edge of feasibility, and thus the distribution of distances, which completely characterizes both shape and size of the feasibility domain. Let us first see what such a question becomes in the abstract -but simpler- case of triangles drawn in the plane, which are the simplest polytopes.

All we need for now is basic trigonometry. If we choose a point in a given triangle, what is the probability that it lies at a distance larger that $z$ from an edge? To answer we need the area $\mathcal{A}_z$ of a smaller triangle whose edges lie exactly at distance $z$ from the borders of the original one (see Fig. 2). We leave it as an entertaining exercise to show that

$$\frac{\mathcal{A} - \mathcal{A}_z}{\mathcal{A}} = 1 - \left(1 - \frac{z}{2\mathcal{A}/\mathcal{P}}\right)^2$$

where $\mathcal{P}$ is the perimeter of the original triangle. The ratio $2\mathcal{A}/\mathcal{P}$ has the dimension of a length, which we can call $D_*$. It is the radius of the largest disc that the triangle can contain. Its center is called the incenter. Thus the proportion $p(z)$ of points that are within distance $z$ from an edge is entirely determined by the ratio $z/D_*$, since

$$\mathbb{P}(\text{ distance } \leq z) = 1 - \left(1 - \frac{z}{D_*}\right)^2$$

### 3 Ecological theory

To build the ecological theory we need a notion of distance, defined in the space of growth rate vectors. For that we take a perturbative perspective. A perturbation $\delta r$ of growth rates leads to a shift $\delta N = A^{-1}\delta r$. If we focus on a given species $i$, this shift reads

$$\delta N_i = \left\langle v^{(i)}, \delta r \right\rangle$$

where $v^{(i)}$ denotes the $i$-th row of the inverse matrix $A^{-1}$, which encodes that species sensitivity to environmental perturbations. Coexistence is lost as soon as one species goes extinct. This implies that $\left\langle v^{(i)}, \delta r \right\rangle = -N_i(r)$ for some species $i$. The smallest intensity necessary, where intensity is measured as

$$\text{intensity } = \sqrt{S}\frac{\|\delta r\|}{\|r\|}$$

gives us a notion of distance $d$ to the edge of coexistence:

$$\begin{aligned} d &= \min \{ \text{ intensity } \mid \delta N_i = -N_i \text{ for some } i \} \\ &= \min \left\{ \text{ intensity } \mid \left| \left\langle v^{(i)}, \delta r \right\rangle \right| = N_i \text{ for some } i \right\} \\ &= \min \left\{ \text{ intensity } \mid \left\| v^{(i)} \right\| \times \|\delta r\| = N_i \text{ for some } i \right\} \\ &= \min_i \frac{\sqrt{S}}{\|r\|} \frac{N_i}{\left\| v^{(i)} \right\|} \end{aligned}$$

In the third line we used Cauchy-Schwartz inequality, more precisely we assumed the worst perturbation $\delta r$ in the sense that it saturates Cauchy-Schwarz's inequality for one species $i$. In the last line we used the expression for intensity. The definition of intensity chosen connects with the Gaussian integral viewpoint as it amounts to enforce $\|r\|^2 = S$, which is the expected norm of growth rate vectors whose components $r_i$ are drawn from $S$ independent standard Gaussian distributions. We can now easily compute the incenter $r^*$ and its radius $D_*$. If we write

$$w_i = \left\| v^{(i)} \right\|$$

representing a species maximal sensitivity, the point that lies at same distance $D_*$ from all edges of feasibility satisfies

$$\frac{\sqrt{S}}{\|r^*\|} \times \frac{N_i\left(r^*\right)}{w_i} = D_*, \forall i$$

thus, if we define the vector of species maximal sensitivities $w = (w_i)$, we have[1]

[1] If we denote $e^{(i)}, i = 1, ..., S$ the canonical vectors of $\mathbb{R}^S$ then we have that

$$w_i^2 = \left\langle A^{-\top} e_i, A^{-\top} e_i \right\rangle = \left\langle e_i, \left(A^\top A\right)^{-1} e_i \right\rangle$$

$$N\left(r^{*}\right)=A^{-1} r^{*}=\frac{\left\|r^{*}\right\|}{\sqrt{S}} D_{*} w \Leftrightarrow \sqrt{S} \frac{r^{*}}{\left\|r^{*}\right\|}=D_{*} A w$$

Note that $r^{*}$ is indeed contained in the feasibility domain because, for all species, $N_{i}\left(r^{*}\right)=D_{*} w_{i}>0$ . All in all this gives us a remarkably simple expression for the maximal distance $D_{*}$ :

$$D_{*}=\frac{\sqrt{S}}{\|A w\|}$$

With this normalization, $D_{*}=1$ in the absence of interactions. Note that

$$r_{i}^{*}=\sum_{j} A_{i j} w_{j}=\sum_{j} A_{i j}\left(A^{\top} A\right)_{j j}^{-1 / 2}$$

is a measure of how effectively hostile, the community -as a whole- is to species $i$. If $r_{i}^{*}=1$, the community has a neutral effect, equal to that of the species on its own. If it is larger than 1 , the community is overall hostile for that species. If it is less than 1 , the community is benevolent for that species.

To understand edge effects, we now make a slight change of perspective, and go back to a probabilistic approach. Instead of asking growth rate vectors to be strictly normalized, we ask that they are normalized on average, meaning that over their distribution, $\mathbb{E}\|r\|^{2}=S$. The two conditions becomes effectively equivalent at large $S$. That being said, recall that $\Phi_{A^{\top} A}(-x)$ is the Gaussian cumulative function associated to $A$, and whose argument $x=\left(x_{i}\right)$ encodes the lower bounds of integration. The probability that growth rate vectors are feasible and further than a distance $z$ amounts to computing the volume of the set of abundances such that $\left\{N_{i} \geq z w_{i}\right\}$ which is $\Phi_{A^{\top} A}(-z w)$, while the conditional probability is $\frac{\Phi_{A^{\top} A}(-z w)}{\Phi_{A^{\top} A}(0)}$. Thus the proportion $p(z)=\mathbb{P}(d \leq z)$ of points that are within distance $z$ (measured as minimal perturbation intensity) from the edge of coexistence is

$$\mathbb{P}(d \leq z)=1-\frac{\Phi_{A^{\top} A}(-z w)}{\Phi_{A^{\top} A}(0)}$$

Now, using the properties of Gaussian cumulative functions allows us to derive the initial slope of $\mathbb{P}(d \leq z)$ :

$$\frac{d}{d z}\Big|_{z=0} \mathbb{P}(d \leq z)=S \sqrt{\frac{2}{\pi}} \times\left(\frac{1}{S} \sum_{i=1}^{S} w_{i} \sqrt{\frac{\left|A^{\top} A\right|}{\left|\left(A^{\top} A\right)_{/ i}\right|}} \frac{\Omega / i}{\Omega}\right)$$

where $\left(A^{\top} A\right)_{/ i}$ denotes the $(S-1) \times(S-1)$ matrix constructed by removing the $i$-th column and row from the original matrix $A^{\top} A$, and where we identified

which shows that so $w_{i}$ is the square root of the corresponding diagonal term of the positive definite matrix $\left(A^{\top} A\right)^{-1 / 2}$.

the probability ratios $\Omega = 2^S \Phi_{A^\top A}(0)$ and $\Omega_{/i} = 2^{S-1}\Phi_{(A^\top A)_{/i}}(0)$. We deduce that in the absence of interactions $p'(0) = S\sqrt{\frac{2}{\pi}}$. We have therefore identified a characteristic distance $D$ that controls for pure dimensionality effects as

$$1/D = \frac{1}{S}\sum_{i=1}^{S} w_i \sqrt{\frac{|A^\top A|}{\left|(A^\top A)_{/i}\right|}\frac{\Omega/i}{\Omega}}$$

Thus we have the initial slope (and value, trivially 0) of $\mathbb{P}(d \leq z)$. We also have that $\mathbb{P}(d \leq D_*) = 1$. So we can get an ansatz that has the characteristics of a cumulative function, mimics the formula for usual triangles, while accounting for dimensionality effects:

$$\mathbb{P}(d \leq z) \approx 1 - \left(1 - \frac{z}{D_*}\right)^{S\sqrt{\frac{2}{\pi}}\frac{D_*}{D}}$$

One can readily check that the initial slope of the expression on the right hand side is indeed $S\sqrt{\frac{2}{\pi}}/D$, that it is otherwise always increasing and reaches 1 for $z = D_*$. This Ansatz is only an approximation, but captures the most salient features of the distribution of distances to the edge of coexistence, and importantly shows fow to relate this distribution to specific features of the interaction matrix $A$.

### 4 Analytical expression in the mean field case

Consider the simplest non trivial interaction matrix

$$A = \begin{pmatrix} 1 & \mu/S & \dots & \mu/S \\ \mu/S & 1 & & \vdots \\ \vdots & & \ddots & \mu/S \\ \mu/S & \dots & \mu/S & 1 \end{pmatrix}; -1 < \mu < S$$

We introduce some useful parameters:

$$\hat{\mu} = \frac{\mu}{1 - \mu/S}; \text{ and } a = \hat{\mu}(2 + \hat{\mu}) > -\frac{S}{S+1} > -1$$

and note that $1 + a = (1 + \hat{\mu})^2$. With these notations the relative volume of the feasibility domain is

$$\Omega \equiv \Omega(S, a) = \sqrt{1+a}\sqrt{\frac{2}{\pi}}\int_{\mathbb{R}_+^S} \exp\left(-\frac{\|x\|^2}{2} - \frac{a}{S}\frac{X^2}{2}\right) d^S x; X = \sum_{i=1}^{S} x_i$$

With the convention that $\sqrt{-1} = i$ we will show that

$$\Omega(S,a) = \frac{1}{\sqrt{2\pi}} \int_{\mathbb{R}} e^{-y^2/2} \operatorname{erfc}\left(i\sqrt{\frac{\lambda(a)}{S}}y\right)^S dy; \lambda(a) = \frac{1}{2}\frac{a}{1+a}$$

where $\operatorname{erfc}(z)$ is the complementary error function. The above can be approximated, as long as $\frac{\lambda}{S}$ is small enough, as

$$\Omega(S,a) \approx \begin{cases} \exp\left(-\frac{S}{\pi}\frac{a}{1+a}\right) & \text{if } a < 0 \text{ (mutualism)} \\ \sqrt{\frac{1+a}{1+Ca}} \exp\left(-\frac{S}{\pi}\frac{a}{1+Ca}\right) & \text{if } a \geq 0 \text{ (competition)} \end{cases} ; \text{ where } C = \frac{\pi - 2}{\pi} \approx 0.36$$

furthermore, we will show that

$$D_* = \frac{1}{\sqrt{1+a-a/S}}; D = \frac{\Omega(S,a)}{\Omega(S-1, a-a/S)} \approx \begin{cases} \exp\left(-\frac{a}{\pi}\frac{2+a}{(1+a)^2}\right) & \text{if } a < 0 \text{ (mutualism)} \\ \exp\left(-\frac{a}{\pi}\frac{2+Ca}{(1+Ca)^2}\right) & \text{if } a \geq 0 \text{ (competition)} \end{cases}$$

This shows, in particular, that

$$\Omega^{2/S} \approx D$$

#### Preparation of $A^\top A$

Let $P_1 = \frac{|1\rangle\langle 1|}{S}$ be the orthonormal projector on the direction spanned by the vector $(1, 1..., 1)^\top$, and $P_1^\perp$ the complementary projector. The spectral decomposition of the mean field interaction matrix takes the form

$$A = (1 - \mu/S)\left[P_1^\perp + (1+\hat{\mu})P_1\right]$$

so we can deduce that $|A| = (1-\mu/S)^S(1+\hat{\mu})$ and also

$$A^\top A = (1-\mu/S)^2\left[P_1^\perp + (1+\hat{\mu})^2 P_1\right] = (1-\mu/S)^2\left[\mathbb{I} + \hat{\mu}(2+\hat{\mu})\frac{|1\rangle\langle 1|}{S}\right]$$

#### Competitive case:

Use the identity

$$\exp\left(-\frac{a}{S}\frac{X^2}{2}\right) = \sqrt{\frac{S}{2\pi a}} \int \exp\left(-\frac{1}{2}\left(S\frac{y^2}{a} + 2iXy\right)\right) dy$$

to show that

$$\Omega = \sqrt{\frac{S}{2\pi}} \int_{\mathbb{R}} \sqrt{\frac{1+a}{a}} dy \exp\left(-\frac{S}{2}y^2\frac{1+a}{a}\right)\left(\sqrt{\frac{2}{\pi}}\int_0^\infty \exp\left(-\frac{(x+iy)^2}{2}\right)dx\right)^S$$

If we set $\lambda = \lambda(a) = \frac{1}{2}\frac{a}{1+a}$ after a change of variables we get

$$\Omega = \sqrt{\frac{S}{2\pi}} \int_{\mathbb{R}} dy \exp\left(-\frac{S}{2}y^2\right) \left(\sqrt{\frac{2}{\pi}} \int_0^{\infty} \exp\left(-\frac{(x+iy\sqrt{2\lambda})^2}{2}\right) dx\right)^S$$

By contour integration in the complex plane we get

$$\sqrt{\frac{2}{\pi}} \int_{i\sqrt{2\lambda}y}^{i\sqrt{2\lambda}y+\infty} \exp\left(-\frac{x^2}{2}\right) dx = 1 - \sqrt{\frac{2}{\pi}} \int_0^{i\sqrt{2\lambda}y} \exp\left(-\frac{x^2}{2}\right) dx$$

and recognize the error function

$$\sqrt{\frac{2}{\pi}} \int_0^z \exp\left(-\frac{x^2}{2}\right) dx = \operatorname{erf}\left(\frac{z}{\sqrt{2}}\right)$$

as well as the imaginary error function

$$\operatorname{erfi}(y) = -i \operatorname{erf}(iy)$$

We finally get to an exact expression:

$$\Omega(S,a) = \frac{1}{\sqrt{2\pi}} \int_{\mathbb{R}} dy \exp\left(-\frac{y^2}{2}\right) \left(1 - i \operatorname{erfi}\left(\sqrt{\frac{\lambda(a)}{S}}y\right)\right)^S ; \lambda(a) = \frac{1}{2}\frac{a}{1+a}$$

To get a simpler but approximate formula, we can linearize the error function near 0 , which leads to

$$-i \operatorname{erfi}\left(\sqrt{\frac{\lambda}{S}}y\right) \approx -2i\sqrt{\frac{\lambda}{\pi S}}y$$

now $1 - i$ erfi $\left(\sqrt{\frac{\lambda}{S}}y\right) = \rho e^{i\theta}$ with

$$\rho \approx \sqrt{1 + \frac{4\lambda}{\pi S}y^2}; \theta \approx -2\sqrt{\frac{\lambda}{\pi S}}y - 2\pi k; k \in \mathbb{Z}$$

thus

$$S \log\left(1 - i \operatorname{erfi}\left(\sqrt{\frac{\lambda}{S}}y\right)\right) \approx \frac{S}{2} \log\left(1 + \frac{4\lambda}{\pi S}y^2\right) - i\left(2\sqrt{\frac{\lambda S}{\pi}}y + 2\pi k S\right)$$

$$\approx \frac{2\lambda}{\pi}y^2 - i\left(2\sqrt{\frac{\lambda S}{\pi}}y + 2\pi k S\right)$$

So

$$\Omega \approx \frac{1}{\sqrt{2\pi}} \int_{\mathbb{R}} dy e^{-i2\pi k S} \exp\left(-\frac{1}{2}\left\{y^2\left(1-\frac{4\lambda}{\pi}\right)+4i\sqrt{\frac{\lambda S}{\pi}}y\right\}\right)$$

$$\Omega \approx \exp\left(-S\frac{\frac{2\lambda}{\pi}}{1-\frac{4\lambda}{\pi}}\right)\frac{1}{\sqrt{2\pi}}\int_{\mathbb{R}} dy \exp\left(-\frac{1-\frac{4\lambda}{\pi}}{2}\left\{\left(y+i\frac{2\sqrt{\frac{S\lambda}{\pi}}}{1-\frac{4\lambda}{\pi}}\right)^2\right\}\right)$$

$$\Omega(S,\lambda) \approx \frac{\exp\left(-S\frac{\frac{2\lambda}{\pi}}{1-\frac{4\lambda}{\pi}}\right)}{\sqrt{1-\frac{4\lambda}{\pi}}} = \sqrt{\frac{1+a}{1+Ca}}\exp\left(-S\frac{a/\pi}{1+Ca}\right)$$

#### Mutualistic case:

Use the identity

$$\exp\left(\frac{|a|}{S}\frac{X^2}{2}\right) = \sqrt{\frac{S}{2\pi|a|}}\int \exp\left(-\frac{1}{2}\left(S\frac{y^2}{|a|}+2Xy\right)\right)dy$$

Using this

$$\Omega = \sqrt{\frac{1+a}{|a|}}\sqrt{\frac{S}{2\pi}}\int_{\mathbb{R}} dy \exp\left(-\frac{S}{2}\frac{y^2}{|a|}\right)\left(\sqrt{\frac{2}{\pi}}\int_0^\infty \exp\left(-\frac{(x+y)^2}{2}+\frac{y^2}{2}\right)dx\right)^S$$

$$= \sqrt{\frac{1+a}{|a|}}\sqrt{\frac{S}{2\pi}}\int_{\mathbb{R}} dy \exp\left(-\frac{S}{2}y^2\frac{1+a}{|a|}\right)\left(\sqrt{\frac{2}{\pi}}\int_0^\infty \exp\left(-\frac{(x+y)^2}{2}\right)dx\right)^S$$

if we set $\lambda = \frac{1}{2}\frac{|a|}{1+a}$ then

$$\Omega = \sqrt{\frac{S}{2\pi}}\int dy \exp\left(-\frac{S}{2}y^2\right)\left(\sqrt{\frac{2}{\pi}}\int_{\sqrt{2\lambda}y}^{+\infty}\exp\left(-\frac{x^2}{2}\right)dx\right)^S$$

We have that

$$\sqrt{\frac{2}{\pi}}\int_{\sqrt{2\lambda}y}^{+\infty}\exp\left(-\frac{x^2}{2}\right)dx = 1-\sqrt{\frac{2}{\pi}}\int_0^{\sqrt{2\lambda}y}\exp\left(-\frac{x^2}{2}\right)dx$$

We can recognize the error function

$$\sqrt{\frac{2}{\pi}}\int_0^z \exp\left(-\frac{x^2}{2}\right)dx = \operatorname{erf}\left(\frac{z}{\sqrt{2}}\right)$$

and we finally get to an exact expression:

$$\Omega(S,a) = \frac{1}{\sqrt{2\pi}}\int_{\mathbb{R}} dy \exp\left(-\frac{y^2}{2}\right)\left(1-\operatorname{erf}\left(\sqrt{\frac{\lambda(a)}{S}}y\right)\right)^S ; \lambda = \frac{1}{2}\frac{|a|}{1+a}$$

To get a simpler but approximate formula, we can linearize the error function near 0

$$-\operatorname{erf}\left(\sqrt{\frac{\lambda}{S}}y\right) \approx -2\sqrt{\frac{\lambda}{\pi S}}y$$

thus

$$S\log\left(1-\operatorname{erf}\left(\sqrt{\frac{\lambda}{S}}y\right)\right) \approx -2\sqrt{\frac{\lambda S}{\pi}}y$$

So

$$\Omega \approx \frac{\exp\left(S\frac{2\lambda}{\pi}\right)}{\sqrt{2\pi}}\int_{\mathbb{R}} dy \exp\left(-\frac{1}{2}\left\{\left(y+2\sqrt{\frac{\lambda S}{\pi}}\right)^2\right\}\right) = \exp\left(S\frac{2\lambda}{\pi}\right) = \exp\left(-\frac{S}{\pi}\frac{a}{1+a}\right)$$

#### Computing $D$

Start from

$$\langle x, A^\top A_{/i}x\rangle_{S-1} = (1-\mu/S)^2\left(\|x\|^2_{S-1} + \left(\frac{S-1}{S}\right)a\frac{X^2_{S-1}}{S-1}\right)$$

from which we deduce the spectral decomposition of $A^\top A_{/i}$ :

$$\begin{aligned} A^\top A_{/i} &= (1-\mu/S)^2\left(\mathbb{I}_{S-1} + \left(\frac{S-1}{S}\right)a\frac{|1\rangle\langle 1|_{S-1}}{S-1}\right) \\ &= (1-\mu/S)^2\left(P_1^\perp + (1+a(1-1/S))P_1^\perp\right) \end{aligned}$$

Thus

$$\sqrt{\left|(A^\top A)_{/i}\right|} = (1-\mu/S)^{(S-1)}\sqrt{1+a(1-1/S)}$$

and so:

$$\Omega_{/i} = \sqrt{(1+a(1-1/S)}\sqrt{\frac{2}{\pi}}^{S-1}\int_{\mathbb{R}_+^{S-1}}\exp\left(-\frac{\|x\|^2}{2} - (1-1/S)a\frac{X^2}{S-1}\right)d^{S-1}x$$

So the same expression as $\Omega$ but with $S \to S-1$ and $a \to a_{/i} = a - a/S$ so

$$\Omega_{/i} = \Omega\left(S-1, a_{/i}\right)$$

and then

$$D = \frac{\Omega}{\Omega/i} \approx \exp\left(-\frac{a/\pi}{1+Ca}\right)\exp\left(\frac{S}{\pi}\left(\frac{a/i}{1+Ca_{/i}} - \frac{a}{1+Ca}\right)\right)$$
$$\frac{a/i}{1+Ca_{/i}} = \frac{a}{1+Ca}\frac{1-1/S}{1-\frac{1}{S}\frac{Ca}{1+Ca}} \approx \frac{a}{1+Ca}\left(1-\frac{1/S}{1+Ca}\right)$$

so

$$S\left(\frac{a/i}{1+Ca/i} - \frac{a}{1+Ca}\right) \approx -\frac{a}{(1+Ca)^2}$$

and therefore

$$D \approx \exp\left(-\frac{a/\pi}{1+Ca}\left(\frac{2+Ca}{1+Ca}\right)\right) \approx \exp\left(-\frac{a}{\pi}\frac{2+Ca}{(1+Ca)^2}\right)$$

For mutualism the expression is simpler

$$\exp\left(-\frac{1}{\pi}\frac{a_{/i}}{1+a_{/i}}\right)\exp\left(-\frac{S}{\pi}\left(\frac{a}{1+a} - \frac{a_{/i}}{1+a_{/i}}\right)\right)$$
$$\frac{a_{/i}}{1+a_{/i}} = \frac{a}{1+a}\frac{1-1/S}{1-\frac{1}{S}\frac{a}{1+a}} \approx \frac{a}{1+a}\left(1-\frac{1}{S}\frac{1}{1+a}\right)$$
$$D \approx \exp\left(-\frac{a}{\pi}\frac{2+a}{(1+a)^2}\right)$$
